# Eigenmode decomposition of asymmetries in whole-brain effective connectivity reveals multiscale hierarchical dynamics

**DOI:** 10.1101/2025.11.16.688716

**Authors:** Giorgia Baron, Giacomo Baggio, Massimiliano Facca, Danilo Benozzo, Alessandro Chiuso, Sandro Zampieri, Alessandra Bertoldo

## Abstract

The human brain is a complex, hierarchical system operating far from equilibrium, yet the mechanisms linking its directed network architecture to temporal irreversibility remain largely unknown. While prior studies have revealed functional and structural hierarchies, no framework has captured the hierarchical organization of the brain’s directed network in relation to its non-equilibrium dynamics. Here, we present the first large-scale eigendecomposition of whole-brain effective connectivity (EC), estimated from resting-state fMRI using sparse Dynamic Causal Modeling. We isolate the irreversible component of EC, which encodes the directionality of information flow and forms a comprehensive hierarchical network of forward and backward interactions. This hierarchy strengthens in brain states approaching criticality, where slow, oscillatory modes dominate and reflects the influence of long-range anatomical connections that scaffold whole-brain information flow. Our decomposition provides the first multiscale quantification of temporal irreversibility, revealing dual counterpropagating streams along the unimodal–transmodal axis operating at distinct frequencies. Crucially, these hierarchical dynamics carry robust, individual-specific signatures, with unimodal networks contributing disproportionately to subject identifiability. Altogether, this work delivers the first dynamic, whole-brain characterization of effective connectivity-derived hierarchies across spatiotemporal scales and individuals, offering a unified framework for studying brain hierarchy, non-equilibrium dynamics, and subject identifiability.

## Introduction

The human brain at rest engages in complex, synchronized interactions across distributed regions, generating dynamic neural activity that is structured both spatially and temporally. Unlike digital systems, the brain operates with relatively slow transmission speeds yet consistently outperforms artificial systems in complex cognitive tasks. This apparent paradox is explained by the brain’s hierarchical architecture, where sensory inputs are transformed stepwise into abstract representations through nested, recursive circuits operating across multiple spatial and temporal scales ^1^. How such emergent dynamics arise from inter-regional interactions—rather than from independent activity alone—remains an open question ^2^.

Previous studies have described global functional harmonic modes, or gradients, that capture principal axes of cortical organization and reveal functional hierarchies embedded in large-scale connectivity ^3,4^. Complementary work has shown that structural harmonics link anatomical and functional organization, suggesting that structural modes can shape large-scale brain dynamics ^5–8^. However, connectome harmonics derived from the graph Laplacian implicitly assume symmetry, overlooking the well-established asymmetries between forward and backward connections^5^. Moreover, undirected models yield time-reversible trajectories, which cannot account for the irreversibility that characterizes empirical brain dynamics ^9^. Addressing these limitations requires dynamic approaches that explicitly incorporate directionality, causality, and hierarchical organization across scales.

Here, we advance this line of research by introducing a mechanistic framework based on sparse Dynamic Causal Modeling (sDCM). Unlike conventional FC-based approaches, the DCM framework estimates directed effective connectivity (EC) and provides a generative model of large-scale brain dynamics ^10^. Using resting-state fMRI data from 144 subjects across two runs of the Human Connectome Project ^11^, we estimated subject-level EC matrices and applied spectral decomposition to their associated complex eigenmodes.

Our study represents the first large-scale eigendecomposition of effective connectivity. This procedure offers a principled tool for characterizing spatiotemporal patterns of neural activity across timescales, each defined by distinct kinetic energy levels ^12^. We show that the irreversible component of EC encodes the directionality of information flow, forming a hierarchical chain of forward and backward interactions. This organization becomes more pronounced near criticality and aligns with structural constraints. Specifically, as the hierarchy expands from local to broader scales, we observe that its functional links show an increasing dominance of anatomical connections with longer fiber lengths, confirming previous hypotheses regarding the critical role of long-range communication ^13^. Furthermore, our decomposition provides the first multiscale quantification of temporal irreversibility, revealing a dual whole-brain hierarchical stream that traverses the unimodal–transmodal axis in opposite directions and at distinct frequencies.

Our work represents the first extensive attempt to link the hierarchical structure of the whole brain’s directed network with its non-equilibrium dynamics across multiple spatiotemporal scales. In doing so, we provide a whole-brain causal and mechanistic account of both the thermodynamic arrow of time and prior observations of traveling waves, which have thus far been largely restricted to surface EEG or phenomenological fMRI models ^10,14–20^, leaving unresolved the interacting network mechanisms through which non-equilibrium dynamics emerge at the macroscopic scale ^10^. Furthermore, by employing a DCM-based, vascular-free modeling strategy, our approach overcomes a key limitation of propagation analyses based on BOLD signals, which are often confounded by vascular and physiological processes ^21^. In this way, we advance from descriptive mapping toward a mechanistic, multiscale characterization of irreversibility that establishes the hierarchical organization of spatiotemporal brain dynamics.

Finally, we show that these hierarchical spatiotemporal patterns also encode individual-specific signatures, extending brain fingerprinting approaches that have primarily relied on undirected functional connectivity ^22–24^. While earlier studies emphasized reproducible zero-lag correlations in transmodal regions, our results highlight the importance of directed contributions from unimodal regions, which provide crucial subject-specific information particularly near criticality, where temporal irreversibility is maximal.

Altogether, this work delivers the first dynamic, whole-brain characterization of effective connectivity–derived hierarchies across spatiotemporal scales and individuals, offering a unified framework for studying brain hierarchy, non-equilibrium dynamics, and subject identifiability.

## Results

### Effective connectivity eigendecomposition reveals energy-based irreversible regimes in the complex eigenspace

From a dynamical systems theory perspective, EC matrices capture the intrinsic temporal architecture of neuronal dynamics (Fig. 1A, left). This structure can be analyzed through the spectral decomposition of each individual EC matrix, interpreted as a linear operator whose eigenvalues and eigenvectors describe the system’s intrinsic modes of evolution ^25^(see Equation (1)).

**Figure 1.**
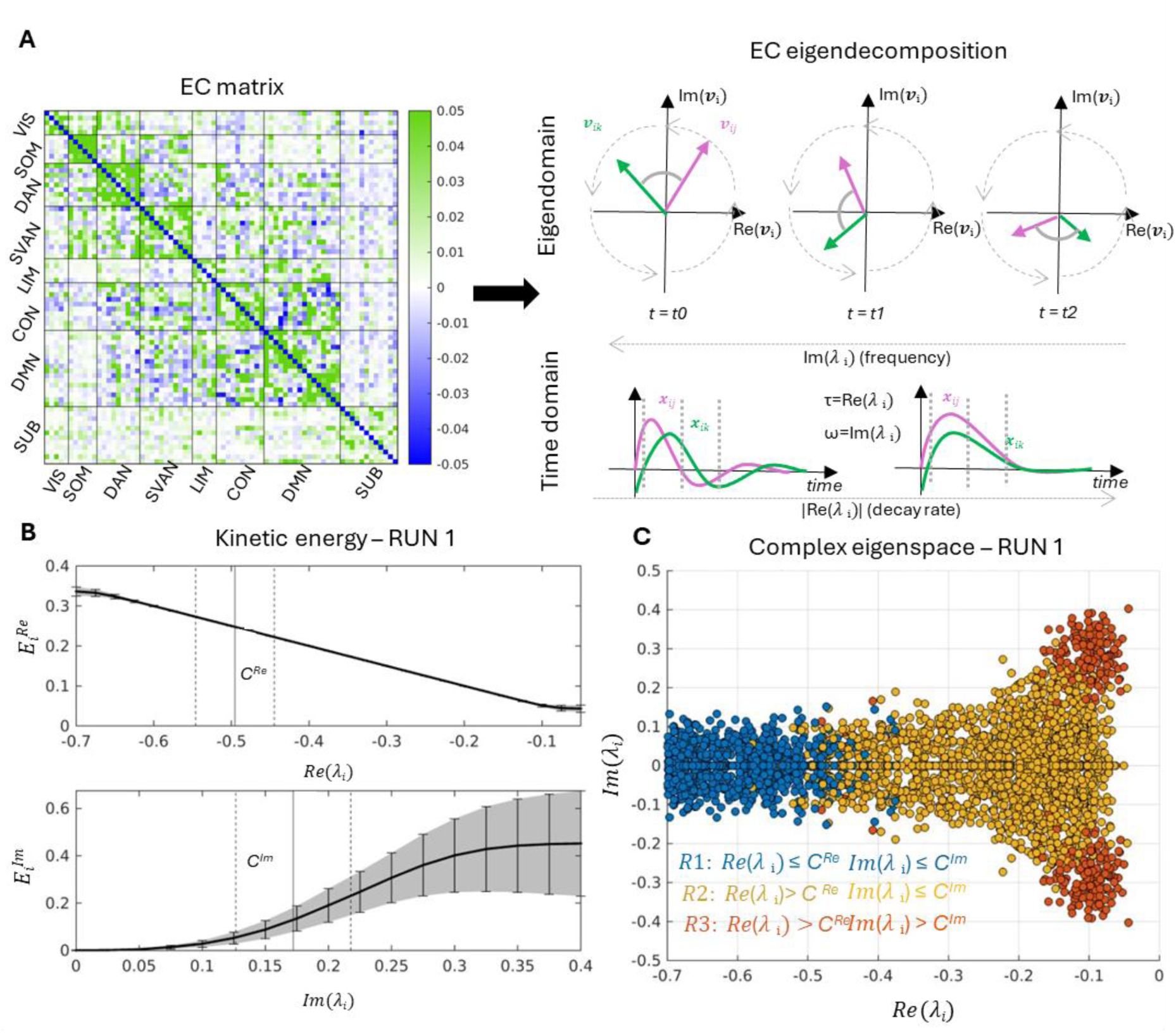
**A)** Example effective connectivity (EC) matrix estimated with sparse DCM (left). The right panel illustrates two forms of EC decomposition. Spectral decomposition is shown at three consecutive time points (t₀, t₁, t₂): dynamics along a single eigenvector dimension can be represented as a rotation in the complex plane, where each eigenvector entry (e.g., *j* and *k*) corresponds to a brain node. For node *j,* the magnitude ∣ *v*_*ij*_ ∣ reflects how strongly that region contributes to the corresponding mode, while the phase *ϕ*_*ij*_ specifies its relative timing within that mode. The oscillation frequency is determined by the eigenvalue’s imaginary component, while the real component encodes decay. For each region signal *x*_*j*_ projected on the mode *i* (i.e., *x*_*ij*_), this relationship translates in the temporal domain as a decaying oscillatory signal (see equation (2)), with dashed grey lines marking the three time points. Increasing real λ leads to faster decay, while increasing imaginary λ corresponds to higher oscillation frequency. **B)** Group-level plots of the real (top) and imaginary (bottom) components of kinetic energy across the eigenspectrum (solid line = mean, shaded area = standard deviation) in Run1. Cut-offs are indicated (solid line = mean, dashed line = CIs). Because the number of modes varied across subjects, individual curves were interpolated within the displayed range to generate the group representation. **C)** Scatterplot of complex eigenvalues across all subjects, shown in the complex plane in Run1. As scales progress from left to right, real eigenvalues converge toward zero. Colors indicate energetic range: blue = first, yellow = second, red = third. On average for Run1, R1 encompasses 7.04 modes, R2 15.35 and R3 2.23.

Because EC matrices are asymmetric, their spectral decomposition occurs in a complex vector space, yielding complex eigenvalues and non-orthogonal eigenvectors that define the system’s dynamic modal decomposition ^26^. Each eigenmode, comprising a real or complex-conjugate pair of λᵢ and associated eigenvectors, characterizes activity in terms of both decay rate and oscillatory frequency (Equation (2)). Specifically, the real and imaginary parts of λᵢ determine the dominant axes along which neuronal trajectories evolve, providing a compact description of the brain’s spatiotemporal dynamics ^27^.

Following ^12^, we interpret each EC eigenvalue in terms of its real and imaginary components, which reflect distinct aspects of intrinsic timescales. The real part quantifies the decay or damping of a mode, reflecting its stabilizing effect, whereas the imaginary part captures oscillatory frequency, associated with solenoidal or rotational flows. According to the thermodynamical analogy outlined in ^12^, these correspond to gradient-driven dissipative flows, which counteract stochastic dispersion, and solenoidal flows, which circulate along isoprobability contours without changing the system’s steady-state distribution. In a complexity plot, this manifests as a rotating, decaying trajectory in the complex plane, where each element’s phase and amplitude maps to a different network region, producing oscillatory-decaying signals in the temporal domain (Fig. 1A, right). In the spectral decomposition illustrated at three consecutive time points (*t₀, t₁, t₂*), the activity of the system along a single eigenmode evolves as a rotation and decay in the complex plane. Within this representation, each point in the complex plane encodes the instantaneous state of all regions projected onto that eigenmode. Regions with similar phase values fluctuate in synchrony, whereas systematic phase differences define a direction of propagation from phase-leading (i.e., region *j*) to phase-lagging (i.e., region *k*) regions. The angular velocity of this rotation is determined by the imaginary part of the associated eigenvalue, which sets the oscillation frequency, while the real part governs the rate of decay of the trajectory toward the origin. In the temporal domain, this corresponds to a decaying oscillatory signal (Equation (2)), where larger real parts of the eigenvalue lead to faster damping, and larger imaginary parts produce higher-frequency oscillations.

To quantify the dynamical relevance of each mode, we computed its kinetic energy using Equation (3) ^12^, separating contributions into real (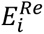) and imaginary (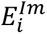) components. 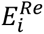 reflects the dissipative, stabilizing energy, while 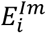 captures the solenoidal, oscillatory energy of each eigenmode. This decomposition allows us to disentangle how each mode contributes to stabilizing versus circulating activity within the system.

Fig. 1B illustrates the spectral evolution of these energy components across the eigenvalue spectrum. A consistent pattern emerged: dissipative energy dominates for highly negative real eigenvalues (fast-decaying modes) and declines near the origin, whereas solenoidal energy increases toward the center, indicating that the slowest, least stable modes exhibit more pronounced oscillatory behavior. This supports theoretical predictions that systems near criticality transition from stable decay to slow, frequency-specific oscillations ^12^. Shaded areas in Fig. 1B show inter-subject variability. Variability is minimal for the dissipative component but increases near the zero real axis for solenoidal energy, suggesting that slow oscillatory regimes are more subject-specific. Occasionally, a few eigenmodes exhibit increased total energy, primarily driven by solenoidal contributions, highlighting the importance of these slow decaying modes in the system’s dynamic repertoire.

Given that each subject’s EC matrix produces a different set of eigenmodes, we implemented a data-driven partition of the eigenspace. Specifically, we divided the area under the curve (AUC) of 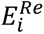 and 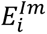 into two equal parts per subject, following a similar procedure to that described in ^7^. Combining the 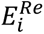 and 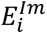 partitions in the complex plane defines four energy-based regimes: R1, with 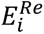 and 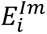 both below the AUC cutoff; R2, with 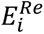 above and 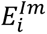 below the AUC cutoff; R3, with both 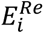 and 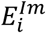 above the AUC cutoff; and R4, with 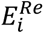 below and 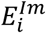 above the AUC cutoff. Notably, R4 was empty across all participants in Run1 (and present in only one subject in Run2; Fig. S8), suggesting that modes with simultaneously high dissipative and solenoidal energy are dynamically unstable or functionally suppressed.

Fig. 1C summarizes the group-level distribution of eigenvalues in the complex plane for Run1. R1 contains eigenvalues with large negative real parts (fast dissipation), while R2 and R3 contains eigenvalues closer to the imaginary axis (slow oscillations). All eigenvalues satisfy *Re(λ)* < 0, ensuring system stability, and appear in complex-conjugate pairs to preserve real-valued dynamics. R2 and R3 eigenvalues are closer to the zero vertical axis, reflecting a small fraction of critically slowing modes exhibiting intrinsic oscillatory behavior at increasing frequencies.

Comparison with previous structural harmonic studies shows that R3 corresponds to the slowest structural eigenvalue range, whereas faster spreading and higher-frequency signaling occurs when transitioning from R3 to R1 ^5,8^. Indeed, higher frequencies encoded by the imaginary component of λᵢ (associated with larger negative eigenvalues) are more strongly attenuated than lower frequencies, consistent with diffusion-like processes. In this context, decay rate (real part) and oscillatory frequency (imaginary part) emerge explicitly from the EC eigendecomposition, providing a direct link between prior results based on structural harmonics, temporal dynamics, and energy distribution.

As outlined earlier, the emergence of complex eigenmodes enables a refined thermodynamical decomposition of effective connectivity, uncovering the spatial organization of information flow that sustains brain dynamics across different timescales. Given our focus on asymmetries in information flow directionality in non-equilibrium systems, we concentrated specifically on the solenoidal component. As ^12^ emphasized, solenoidal coupling introduces directed macroscopic fluxes that break detailed balance—a defining feature of systems operating far from equilibrium. This results in an irreversible “arrow of time” in system dynamics, captured by asymmetries in event flow ^28,29,18^.

In linear models such as sDCM, the solenoidal component can be extracted in closed form (see Equation (3-6) in Supplementary Methods). The antisymmetric part of the effective connectivity matrix, denoted as *S* in the present study, encodes the solenoidal flow driving these asymmetries ^30,31^. Statistically, *S* corresponds to the differential cross-covariance matrix and directly characterizes directional information flow. By construction, if *S_ij_ > 0* (and *S_ji_ < 0* due to skew-symmetry), then node *j* acts as a source and node *i* as a sink, indicating that information is flowing from *j* to *i* ^32^.

This thermodynamic decomposition applies locally at each point in the brain’s state space and enables the spectral projection of the spatial structure of *S* across the three energetic ranges derived from eigendecomposition in the complex domain (see Equation (4)). In this framework, eigenvalues regulate decay and oscillatory rates in the temporal domain, while entries of *S* reveal which regions act as sources or sinks within each range domain, relying on brain dynamics patterns that emerge from the spatial distribution of eigenvector phase and magnitude components (see Fig. 2A, Supplementary Results and Supplementary Fig. S1, S11).

**Figure 2.**
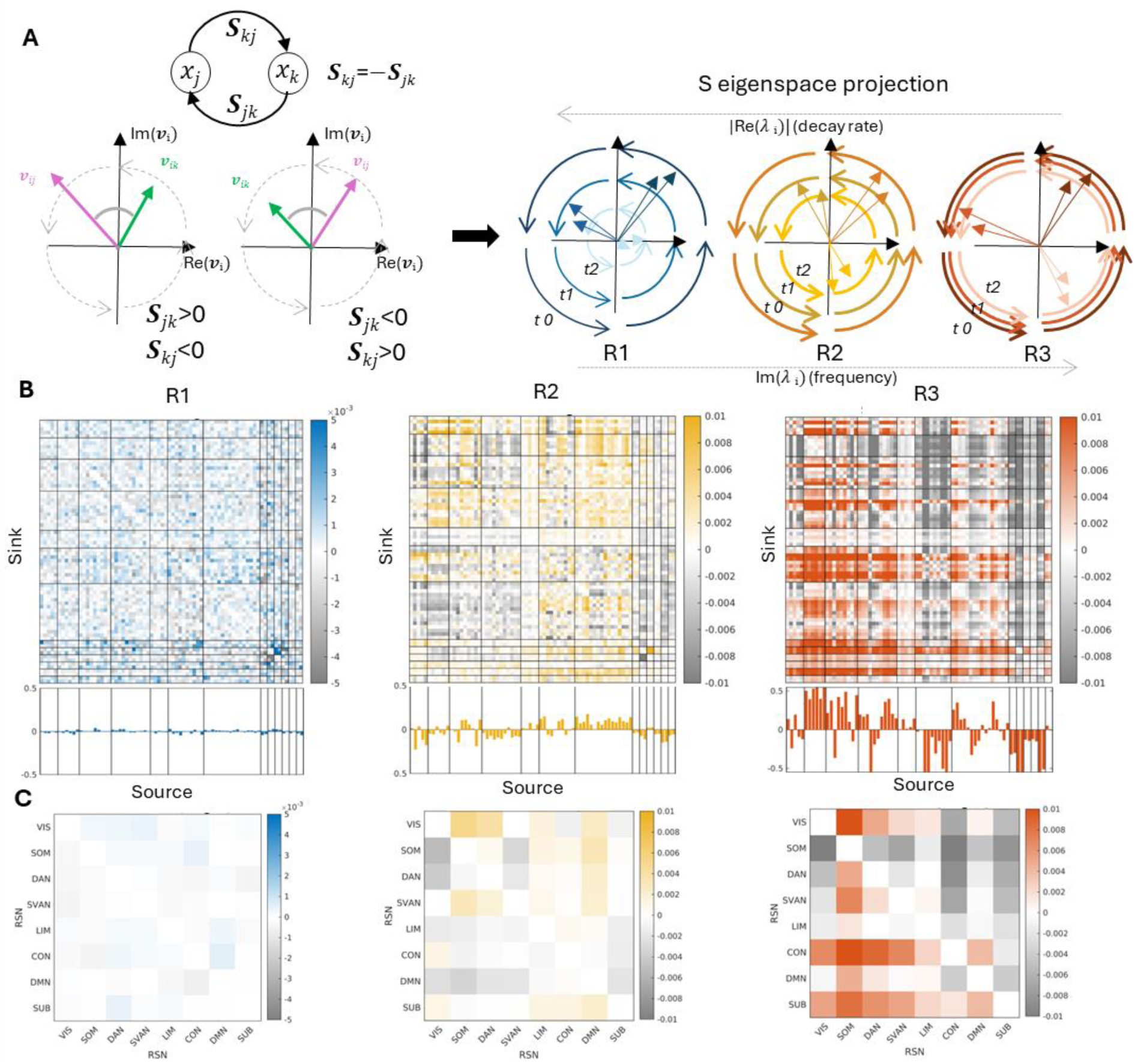
**A)** As shown in Fig. 1A, the dynamics between each pair of nodes can be represented in the complex plane as rotating entries of the complex eigenvector. Accordingly, the corresponding dc-Cov entry in the matrix *S* changes coherently: if *S*_*jk*_ > 0 (and *S*_*kj*_ < 0, by skew-symmetry), node *k* causes node *j*, so *v*_*ik*_ precedes *v*_*ij*_ in the rotating plane; if *S*_*jk*_ < 0 (and *S*_*kj*_ > 0), node *j* causes node *k*, so *v*_*ik*_ follows *v*_*ij*_. When projecting the spatial pattern of S onto the three ranges, differential cross-covariance asymmetries are separated according to the temporal dynamics encoded by the eigenmodes within each range—namely, slower decay rates and increasing oscillation frequencies from R1 to R3. **B)** Group-level average patterns of the differential cross-covariance matrix for the first second and third ranges for Run1. Each individual matrix is normalized by its Frobenius norm, and ROIs are grouped according to cortical and subcortical networks (black lines). Below each matrix, the column-wise strengths are reported: a negative column sum indicates that the node predominantly acts as a *receiver*, while a positive sum indicates that it predominantly acts as a *sender*. **C)** Same matrices as in (B), but averaged across resting-state network (RSN) blocks.

Fig. 2B shows the group-averaged solenoidal matrices at ROI level, along with node strengths, while the same matrices at network level are reported in Fig. 2C. The dC-Cov matrix *S* initially displays a flat spatial pattern, with only a few weak subcortico-cortical links in R1. As the system approaches instability, asymmetric couplings strengthen, peaking in R2 and R3, coinciding with critically slow modes. This escalation marks increasing irreversibility and a departure from detailed balance.

Network differentiation becomes evident in the second range: unimodal systems (VIS,SOM,DAN,SVAN) show negative *S* strength, whereas transmodal networks (LIM,CON, DMN) exhibit positive strength. This suggests that higher-level networks predominantly act as senders, while unimodal networks function as receivers. In the least stable range (R3), this pattern reverses: CON and DMN regions primarily become receivers or weak senders, while somatosensory-motor and attentional areas assume source roles in solenoidal transfer. Subcortical and cerebellar nodes also differentiate more strongly in R2 and R3, increasingly acting as targets for cortical information.

### Large multiscale hierarchies emerge from solenoidal and long-range information flow

The breaking of detailed balance induced by *S* reflects preferential information flow, with asymmetries encoding a hierarchical reorganization of brain dynamics ^31^. A well-defined arrow of time indicates a clear hierarchical organization, with strong breaking of detailed balance and asymmetric information flow that assign distinct computational roles to different areas. Conversely, a weaker arrow of time suggests greater symmetry and a flatter hierarchy of brain organization. In other words, the degree of irreversibility in brain signals provides a direct measure of hierarchy.

The assumption that hierarchies require asymmetry between incoming and outgoing connectivity is equivalent to demanding acyclic information flow, a principle formalized in measures of flow hierarchy ^33^. To identify the main hierarchical streams of whole-brain information flow encoded in *S*, we then derived, for each subject and each range, a directed acyclic graph (DAG). This was achieved through iterative sparsification, progressively eliminating residual cycles (Fig. 3A). The procedure was repeated 100 times, and only connections consistently retained across iterations were used to construct the final individual DAG. Each subject-level DAG was then topologically sorted to establish a unique ranking of brain nodes.

**Figure 3.**
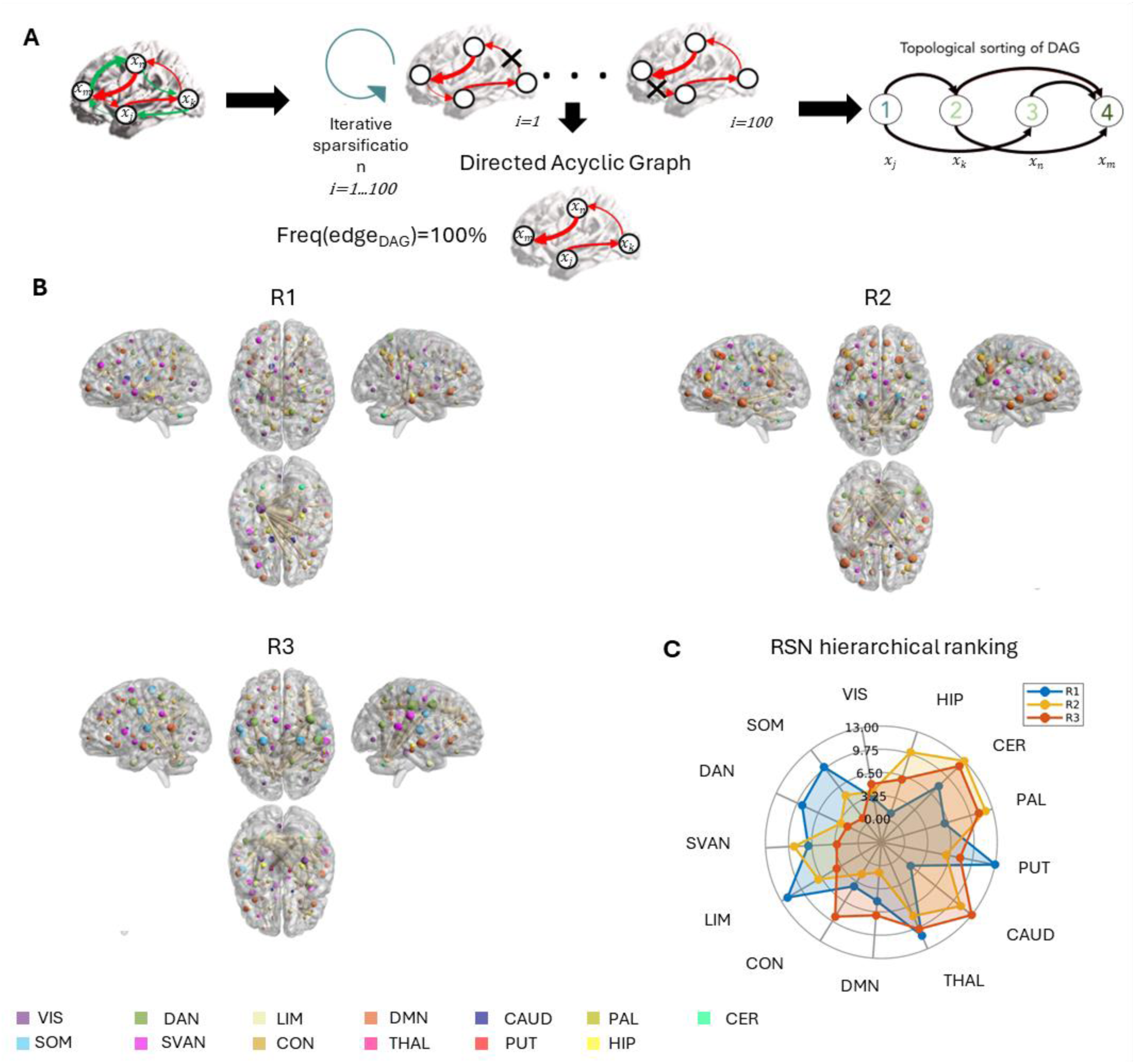
**A)** Workflow for constructing the directed acyclic graph (DAG) of the solenoidal flow. Positive links are extracted, and the minimum number of cycles is removed to obtain a DAG. Because cycle removal involves random choices, the procedure is repeated 100 times for each subject and range. The final DAG is defined by retaining only the links consistently present across all iterations. Finally, regions of interest (ROIs) are ranked hierarchically using topological sorting. **B)** Mean ROI rankings, derived from each ROI’s ranking distribution are shown on a brain rendering to elucidate the spatial distribution of source and target nodes for Run1. Node size reflects the role of the node as a source (larger nodes indicate earlier positions in the hierarchy). Links are displayed if their frequency exceeds the 99^th^ percentile of the DAG frequency matrices (see Supplementary Fig. S5A), with edge thickness proportional to frequency. Node colors correspond to functional networks. **C)** Spider plot showing mean rankings at the cortical and subcortical network level across ranges (lower values = sources; higher values = sinks).

Group-level brain renderings of these hierarchies are shown in Fig. 3B, where increasing node size reflects the prominence of a region as a source. Fig. 3C compares hierarchical orderings across ranges at the network level, while subject- and ROI-level distributions are reported in Supplementary Fig. S2 and S3 (Supplementary Fig. S12 and S13 for Run2). No significant global ordering was detected in the initial range (not significant Jonckheere–Terpstra test, critical α=0.01).

By contrast, R2 and R3 revealed robust complementary spatial hierarchies, with sources and targets interchanging roles as timescales approached instability. More specifically, the cortical ranking observed in R2 was statistically significant (Jonckheere–Terpstra, p<10^−30^ for Run1 and p<0.006 for Run2) and recapitulated the well-established functional gradient from perception and action to higher cognition ^4^. Specifically, asymmetry revealed that information transfer flowed predominantly from higher-order networks (DMN, CON) to attentional and sensory networks (DAN, VIS, SOM, SVAN). Among subcortical nodes, cerebellum, pallidum, caudate, and hippocampus emerged as prominent targets, while thalamus and putamen occupied more central hierarchical positions.

In R3, the cortical hierarchy inverted (Jonckheere–Terpstra test, p<10^−30^ for Run1 and Run2): somatosensory-motor and attentional regions emerged as dominant sources, whereas DMN and CON regions primarily acted as targets. VIS and LIM networks, along with hippocampus, exhibited more balanced roles in temporal directionality. Subcortical and cerebellar nodes consistently occupied lower hierarchical positions in this range.

Since the strength of hierarchy corresponds to the proportion of edges not participating in cycles ^20^, the robustness of hierarchies in R2 and R3 was confirmed by comparison with null DAGs: both the number and weight of removed cycles were consistently lower than expected by chance (Supplementary Fig. S5). Moreover, the stability of the identified acyclic connections was supported by frequency maps across iterations. Comparable results were obtained in Run2 (Supplementary Fig. S15).

Prior work has shown that computation at the network level emerges from critical dynamics producing long-range interactions that give rise to low-dimensional manifolds ^34^. These findings suggest that structural embedding, and in particular the distribution of fiber lengths, plays a key role in supporting functional hierarchies. Motivated by this, we tested whether the hierarchical interactions identified from the solenoidal flow are actually scaffolded by anatomical structure. For each subject, we extracted fiber-length matrices from diffusion data and divided connections into short-, medium-, and long-range classes (see Supplementary Methods and Fig. S4). Comparing the distribution of each class across ranges revealed that R1 was primarily supported by short and medium fibers (Supplementary Fig. S4B and C), whereas R2 and R3 displayed a significantly higher prevalence of long-range fibers (R2 > R3) than R1 (Supplementary Fig. S4D). These results replicated in Run2, with short-range fibers predominating in R1 and long-range fibers in R2 and R3 (Supplementary Fig. S14).

### Hierarchical flows support the fingerprint of unimodal networks

After deriving hierarchies for each subject and each range from the first and second run of LR rs-fMRI (see Supplementary Fig. S8-S15), we assessed their reliability through fingerprinting metrics. We first quantified subject identifiability, the extent to which a subject can be uniquely identified within a group based on connectivity edges. Identifiability was measured using *I_diff_*, the difference between *I_self_* and *I_others_* ^35^, where *I_self_* reflects within-subject similarity across sessions and *I_others_* reflects similarity between different subjects (see Equation (6)). We also computed the success rate (SR) of subject identification as originally proposed by ^22^.

We systematically examined *I_diff_* across subjects, flow patterns, and energetic ranges. The distributions are shown in Fig. 4A. For both *I_diff_*and SR, ANOVA test with subject as a within-subject factor was significant, with post hoc tests revealing range-specific effects. In the initial range (R1), where solenoidal flow has minimal contribution, identifiability values were relatively low. By contrast, identifiability increased across ranges, peaking in R3. This indicates that within-subject similarity surpasses between-subject similarity most strongly in the solenoidal component of the final range (see also *I_self_* and *I_others_* distributions in Supplementary Fig. S7). A similar trend was observed for SR, and R2 and R3 mean *I_diff_* and SR values were significantly different from null distributions according to permutation testing (see details in Methods).

**Figure 4.**
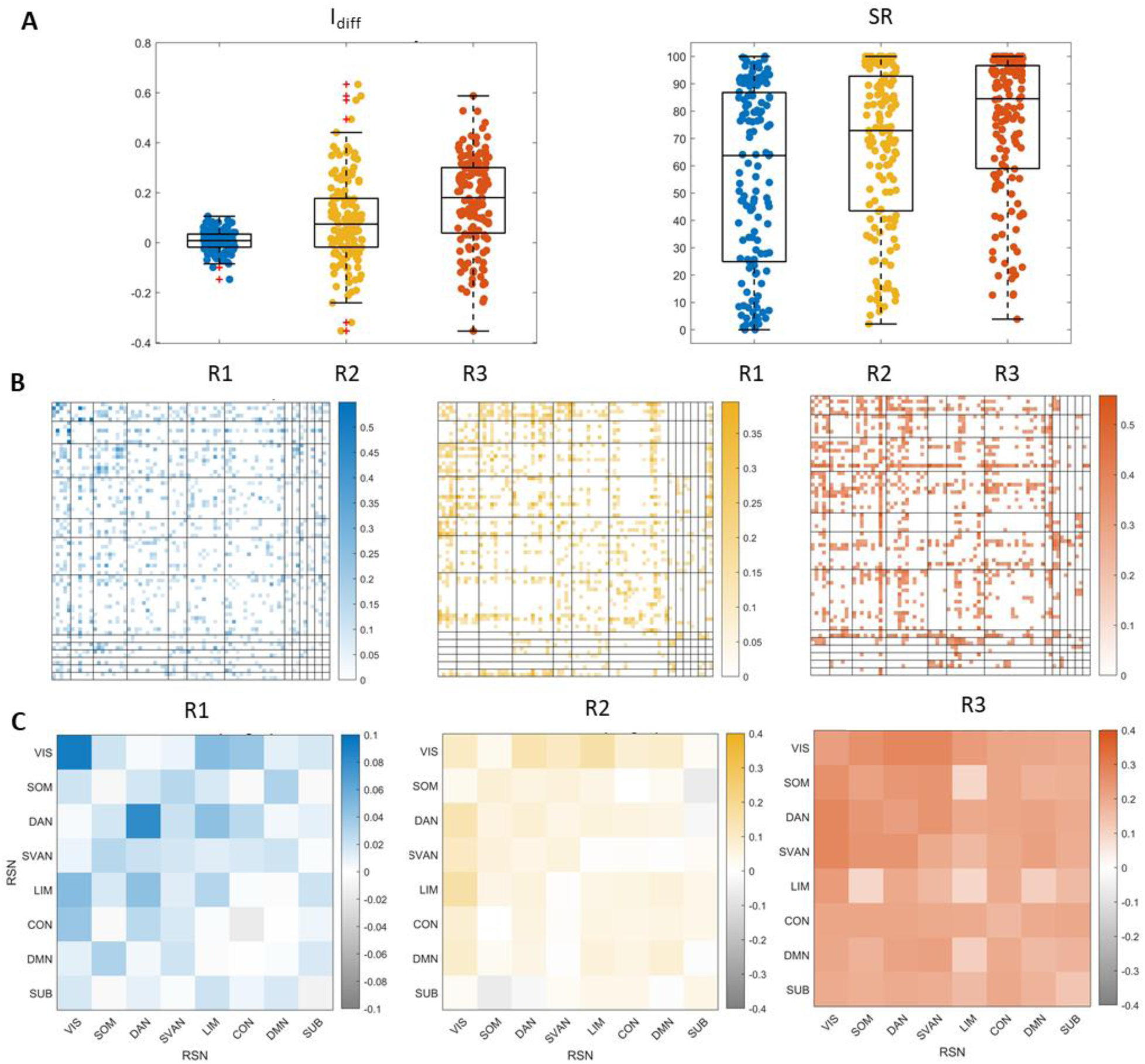
**A)** Boxplots showing the across-subject distributions of differential identifiability (*I_diff_*) and success rate (SR) across the three ranges, with individual values overlaid as jittered points. The boxes display the median and interquartile range of each distribution, with whiskers extending to non-outlier data. Repeated-measures ANOVA revealed a significant main effect of range for both Idiff (F(2, 284) = 50.17, p = 2.19 × 10⁻¹⁹) and SR (F(2, 284) = 16.15, p = 2.28 × 10⁻⁷), and post hoc Tukey–Kramer tests showed that all pairwise differences were significant, with p-values ≤ 0.01. Permutation testing, using a critical α of 0.01, confirmed that mean values in ranges 2 and 3 were significantly greater than zero, while values in range 1 were not. **B)** Edgewise S connectivity fingerprints quantified using intra-class correlation (ICC), displayed for the three ranges and thresholded at the 80th percentile. **C)** Average ICC values within and between resting-state networks (RSNs), reported without thresholding.

To further characterize subject-specific features, we additionally estimated idiosyncrasy using the intraclass correlation coefficient (ICC), a widely used measure of reliability that quantifies how strongly units within the same group resemble each other ^36^. Here, ICC captures the reproducibility of each edge across test–retest (e.g., Run1-Run2) acquisitions, providing a direct measure of fingerprinting value ^35^(Equation (7)).

Edgewise ICC maps, thresholded at the 80^th^ percentile for visualization (Fig. 4B–C), revealed a progressive reorganization of subject-specific features across ranges. In R1, ICC values were relatively low and concentrated within VIS and DAN. In R2, reproducibility increased, with identifiable edges emerging across VIS, motor and attentive RSNs (SOM, DAN, SVAN), transmodal networks (LIM, CON, DMN), and subcortical/cerebellar nuclei. In R3, ICC patterns shifted toward unimodal networks (VIS, SOM, DAN, SVAN), with additional idiosyncratic edges involving thalamus–cortex, caudate–CON, and cerebellum–LIM/CON connections. Importantly, ICC maps did not resemble DAG frequency maps (Supplementary Fig. S5A), showing that the edges most frequently retained in DAGs are not necessarily the most distinctive at the individual level. Interestingly, in R3, within-network unimodal edges were more reproducible, whereas cross-network edges were more frequent in DAGs.

To capture regional contributions, we also computed a nodal fingerprinting score as the sum of ICC strength for each region ^36^, highlighting the spatial organization of regional *S* signatures (Supplementary Fig. S6). From R1 to R3, nodal fingerprinting strength increased and became more widely distributed, indicating stronger node-level specialization. VIS regions consistently displayed high fingerprinting values across ranges. Between R2 and R3, a reconfiguration emerged, with unimodal and transmodal networks alternating in dominance, particularly in the right hemisphere. Subcortical nuclei also gained prominence, with hippocampus emerging in R2, and thalamus, caudate, and cerebellum contributing more strongly in R3.

Together, these results demonstrate that hierarchical solenoidal flows encode robust, individual-specific signatures. Identifiability increases across energetic ranges, peaking in R3, and the spatial distribution of idiosyncratic edges and nodes highlights a shifting balance between unimodal and transmodal contributions to individual fingerprints.

## Discussion

Earlier connectivity studies using graph Laplacians have revealed gradients of cortical organization but are constrained by undirected assumptions, missing causal asymmetries between feedforward and feedback pathways. FC-based fingerprinting has likewise emphasized static correlations in transmodal regions while overlooking directed contributions from unimodal areas. More critically, in neural dynamics, where broken detailed balance consistently emerges, the traditional assumption of undirected connections limits the accuracy of whole-brain models and prevents them from explaining the non-equilibrium nature of empirical data.

Here, we address these limitations by applying spectral decomposition to empirically estimated effective connectivity. This approach captures the solenoidal component of brain dynamics, revealing irreversible, frequency-specific propagation streams along the unimodal–transmodal axis, providing a mechanistic account of hierarchical organization and temporal irreversibility, while also encoding robust subject-specific signatures that show solenoidal patterns are both structured and individualized.

A pivotal novelty provided by this study is the explicit description of whole-brain dynamics, as posited by the dynamic causal modeling framework, as a linear combination of complex eigenmodes. While Laplacian eigendecomposition of connectivity matrices has been widely applied to study structure–function coupling ^5,37,7,24^, its extension to whole-brain EC has been lacking. Because EC is asymmetric, this requires handling complex eigenvalues and eigenvectors ^38^. Their real and imaginary parts encode decay rate and oscillatory frequency, respectively, introducing a dual definition of kinetic energy that captures both dissipative and solenoidal contributions ^12^. This formulation resonates with ^5^, who defined the energy of connectome harmonics from structural connectivity by combining intrinsic energy (*λ*^2^) with activation strength. However, our energy formulation inspired by ^12^ proposed an eigenmode-specific energy grounded in subject-level eigenvalues of EC, thereby embedding inter-individual variability directly into the spectral domain.

Following ^7^, we partitioned each individual eigenspectrum into three energetic ranges (Fig. 1B and C). The first range is dominated by very fast, highly stable modes that dissipate quickly in the temporal domain. In contrast, the second and third ranges reveal increasingly oscillatory behavior, where temporal patterns decay more gradually, with critically slow modes approaching instability from below ^39^. This distinction is clearly visible in the eigenvalue distribution of Fig. 1C, which would remain undetected using symmetric decompositions, and confirms that effective brain networks are characterized by fixed points near instability, where slow oscillatory modes govern long-term dynamics ^38^.

Moving from the real to the complex eigenspace allowed us to capture not only oscillatory dynamics but also phase asynchrony within the spatial domain of brain interactions. Building on ^26^, complex eigenmodes signify traveling waves and net energy flow, while zero-phase eigenvectors correspond to standing waves. This duality is illustrated in the relationship between *S* and complex eigenvectors reported in Supplementary Fig. S1, where phase shifts track solenoidal flow to increasing degree from R1 to R3, and aligns with geophysical insights from ^40^, who argued that intrinsic brain activity reflects a mixture of standing and traveling waves. In their study, three dominant spatiotemporal patterns of spontaneous low-frequency BOLD activity were identified, two of which exhibited traveling-wave properties despite not accounting for inter-participant variability.

This conceptual distinction is evident in *S* patterns (Figs. 2B and C). Standing waves correspond to stationary oscillations with no time-lagged spatial dependence, consistent with the negligible solenoidal contribution in R1. In contrast, traveling waves reflect oscillations with non-zero phase lags, revealed by the solenoidal matrix *S* dominating in R2 and R3. Here, slowly decaying modes shape the cross-covariance structure ^39^, emphasizing the traveling-wave character of propagating activity.

As noted by ^12^, the presence of complex eigenvalues reflects a loss of detailed balance, i.e., entropy production, captured in our framework by asymmetries in S. Node strength in S thus serves as a proxy for induced imbalance ^31^. Examining node strength profiles across RSNs reveals a distinct dissociation: cognitive networks act as moderate senders in R2 with relatively low irreversibility, while unimodal networks dominate in R3, contributing more strongly to entropy production. This agrees with ^29^, who reported that unimodal systems driven by the environment such as SOM exhibit high non-reversibility and reduced flexibility, whereas higher-order networks like DMN and LIM remain more reversible and flexible. However, low irreversibility does not imply absence of directionality. Rather, higher operating frequencies make phase shifts more difficult to detect, producing an apparent standing-wave pattern that can mask more subtle directional propagation ^41^.

As a result, we elucidated how the temporal directionality inherent in *S* can be leveraged to construct a comprehensive whole-brain DAG, revealing the hierarchical structure of information flow at multiple spatiotemporal scales (Fig. 3). In the initial range, hippocampus, caudate, and visual areas emerge as sources projecting toward transmodal, attentive, motor, and other subcortical networks. However, no discernible whole-brain orderings were identified, consistent with the low magnitude of S highlighted in Fig. 2B. Simply put, the observed flat hierarchy is marked by a minimal degree of breaking of detailed balance, signifying largely symmetrical information flow ^29^.

Conversely, a global hierarchy becomes apparent in R2 and R3, consistent with the increasing level of irreversibility ^10^. In range 2, the ordering recalls the transmodal–unimodal gradient ^4^, which describes gradual shifts in functional connectivity profiles from heteromodal regions to primary sensory cortices. Specifically, the hierarchical ordering suggests information flow from DMN, limbic, and executive control motifs generating prediction signals toward primary sensory, attention, and salience motifs. These latter motifs process sensory input and refine internal models through prediction errors ^42^. This is also consistent with more prevalent high-frequency activity expected in cognitive hubs: ^14^ showed that increases in encoded information scale with higher firing rates, and simulations revealed that hubs sustain the highest firing rates and power.

In R3, the global ranking reverses, separating motor and attentive (senders) from cognitive (receivers) networks. This recalls the differentiation between representation and modulation networks, where attentive regions fine-tune the precision of predictions and errors associated with contextual representations in sensory and cognitive areas ^43^. Intriguingly, attentive regions likely sustain broad coordination, integrating ongoing unimodal processing with extended heteromodal operations ^44^. These dynamics suggest that near-critical behavior observed in R3 represents an optimal regime for flexible integration. The separation between somatomotor and visual networks further indicates a segregation of exteroceptive sensory systems, reflecting distinct properties of signals from different sensory surfaces ^42^ and may be linked to subtle but systematic time delays between motor and visual areas during cross-hierarchy propagations ^45^.

Subcortical nodes also display frequency-dependent roles. ^46^ proposed a sender–receiver architecture in hippocampal–cortical dialogue, with higher frequencies supporting sender roles. Our results align with this view: the hippocampus shifts from a lower rank in R1 to a higher receiver role in R2 and R3. Conversely, the cerebellum typically functions as a receiver, consistent with its role in predicting incoming sensory signals and refining the precision of cortical prediction errors ^42^.

Altogether, our findings uncover temporal and spatial directionality underlying a dual hierarchical stream, linking them to the top-down/bottom-up computational processes of predictive coding and the free-energy principle ^43^. This resonates with dual-process theories of cognition ^47^, where fast and slow systems coexist across multiple timescales, and with recent demonstrations of ascending and descending cortical propagations in traveling waves ^21^.

Interestingly, the unveiled hierarchical dynamics also echo electrophysiological evidence. Higher-frequency hierarchy in R2 align with beta-band oscillations reflecting top-down signaling from higher-order to sensory cortices ^48^. By contrast, the slowest hierarchy in R3 near criticality recalls findings that alpha oscillations strongly correlate with critical brain tuning ^49^. The source role of DAN and SVAN networks in R3 is also consistent with the attentional modulation role of alpha waves ^16^. TMS studies corroborate this frequency-dependent hierarchy, showing dominance of alpha in occipital, beta in parietal, and gamma in frontal cortices ^50^. Moreover, ^51^ reported posterior–anterior gradients across theta, alpha, and beta frequencies: alpha mirrored posterior-anterior gradients, while theta and beta showed increasing anterior frequencies, consistent with traveling waves from anterior to posterior cortices. Consistently, ^52^ further identified alpha and beta subnetworks along instrength gradients, with alpha traveling waves coordinating non-zero-lag connectivity. These findings converge with our results, which reveal, for the first time in resting-state fMRI, the hierarchical network mechanisms underlying such propagating activity.

The anatomical underpinnings of these ranges reinforce their functional distinctions. By comparing the distribution of fiber lengths, we found that R1 is supported by a greater proportion of short and medium fibers, while R2 and R3 rely disproportionately on long-range fibers (Fig. S4 and Supplementary Fig. S14).This confirms that in R1 the DAGs fail to converge to a global hierarchy across subjects, reflecting localized, module-specific dynamics ^3^ supported by short fibers. This is consistent with the role of high-frequency eigenmodes in transient, localized spreading processes that support flexible state switching ^53^. However, given the low magnitude of S in Fig. 2B, such interactions are also more unstable and likely more influenced by fMRI noise at high oscillation frequencies.

In contrast, the dominance of long-range fibers in R2 and R3 resonates with evidence that rare long-range connections disproportionately shape network topology to optimize information processing ^1,13,54^. Prior work showed that such exceptions are critical for resting-state networks and enhance turbulence and integration–segregation balance. Crucially, our solenoidal decomposition provides additional insight by revealing the directional contribution of these fibers to global hierarchies. In this way, our framework overcomes the limitations of ^55^, who showed that long-range links enhance hierarchical dynamics but could not infer their causal or directional roles. Here, we demonstrate precisely how these connections encode and drive asymmetric propagation, establishing a mechanistic basis for their contribution to hierarchical brain organization.

Regarding individual fingerprinting, we examined whether solenoidal profiles can uniquely characterize subjects, introducing a novel dimension to inter-individual differences in brain dynamics. At the whole-brain level (Fig. 4A), fingerprinting power increases markedly in the last range when considering time-lag interactions captured by *S*. This indicates that the directional information carried by critically slow and oscillatory eigenmodes contributes substantially to subject identifiability. This trend is in stark contrast with the low identifiability observed in these same ranges when only zero-lag undirected interactions are considered ^8,24^. In those studies, the functional component detached from structure—specifically, the pattern projected onto high-frequency structural harmonics—enabled near-perfect subject identification, but relied mainly on transmodal networks. Similarly, ^56^ found that higher-order heteromodal systems dominate identifiability, whereas unimodal regions contributed little. ^57^ further confirmed that associative systems drive subject identifiability across TRs, while sensory–motor regions are more sensitive to temporal resolution.

Our results challenge this consensus, showing that unimodal and attentive regions also contribute robust fingerprinting power when non-zero-lag interactions are taken into account. This aligns with recent evidence highlighting the role of aperiodic, bursty dynamics in subject differentiation: ^58^ demonstrated that neuronal avalanches, scale-free and aperiodic in nature, propagate across the brain and encode subject-specific signatures, reflecting near-critical activity as a substrate of individual uniqueness.

Turning to the ICC patterns of *S* (Fig. 4B and C), reliability is very low in R1, confirming that high-frequency estimates of *S* are too noisy for stable fingerprinting. In contrast, pronounced specialization emerges in the later ranges, where the most reproducible links involve the visual network in R2 and unimodal (visual, motor, and attentive) ROIs in R3. This is noteworthy, as ^59^ had highlighted the attention network’s metastable regime supporting cognitive control. Here, we show that the directed coupling within such critical regimes also enhances identifiability.

Support for this perspective still comes from EEG/MEG studies that, like our solenoidal framework, quantify connectivity through phase-lag interactions. ^60^ reported robust fingerprinting in the alpha band using phase-based connectivity metrics, particularly in occipito–temporal and occipito–parietal regions, consistent with our R2 findings. Similarly, ^61^ demonstrated that alpha and beta bands yield the highest reliability across both rest and task conditions, again based on phase-lag indices. ^36^ further confirmed strong subject identifiability in these bands, with phase-based measures outperforming others. Their nodal patterns emphasized parieto-occipital and precuneus regions in alpha and somatomotor regions in beta, closely mirroring our observation that visual, motor, and attentive nodes dominate fingerprinting in R2–R3 (Supplementary Fig. S6). Importantly, ^36^ also showed that phase-lag–based fingerprinting in the alpha and beta bands accounted for more than 50% of connectome–cognition covariance, underscoring not only identifiability but also behavioral relevance. This methodological alignment strengthens the validity of interpreting S as a meaningful measure of directed, frequency-specific brain fingerprinting. Although high test–retest reliability does not always translate to behavioral prediction ^62^, robust identifiability provides an upper bound for predictive validity ^35^. Our results indicate that fingerprinting power rooted in unimodal and attentive solenoidal dynamics reflects stable, subject-specific traits and the critical directional processes underlying them.

### Limitations

While this study unveils novel findings regarding connectivity dynamics, some limitations should be acknowledged. First, the use of a limited number of functional parcels, necessary to reduce computational load, constrains our ability to provide a fully nuanced characterization of the information flow patterns derived from effective connectivity. Second, certain factors may impact the dynamic mode decomposition of the EC matrices, highlighting the need for comprehensive sensitivity analyses. Specifically, our analyses are confined to the slow temporal scales accessible with fMRI, which may limit the estimation of phase shifts shaping asymmetric coupling in the first range (R1), dominated by high-frequency oscillations not captured by the low temporal resolution of fMRI. Consequently, the predominance of standing wave patterns observed in R1 may partly reflect this methodological limitation rather than a true absence of directional propagation.

Future research could address these constraints by investigating the hierarchy of information flow at faster temporal scales, leveraging techniques such as MEG or EEG, which could provide a more precise characterization of local hierarchies and allow mapping between the frequency bands observed here and their corresponding neuronal oscillations. Additionally, the heuristic procedure used to remove cycles in DAGs could be refined using more sophisticated approaches, such as algorithms for identifying the minimum feedback arc set ^63^.

Finally, although we employed a linear model to characterize brain dynamics, recent evidence supports the validity of such an approach. ^64^ demonstrated that linear models can be as descriptive as nonlinear ones at the macroscale, while ^65^ showed that linear control model predictions align with nonlinear biophysical simulations for personalized stimulation protocols. Moreover, linear models offer enhanced interpretability compared with their nonlinear counterparts, providing a practical advantage for understanding and modeling whole-brain dynamics ^66^.

### Conclusions

Adopting a thermodynamic perspective, we present a mechanistic explanation for the intricate spatiotemporal dynamics of brain function by analyzing whole-brain effective connectivity through dynamic mode decomposition. By revealing asymmetric coupling through complex eigenspace analysis, our framework provides a unified overview that explains much of the existing literature based on symmetric graphs, while elucidating for the first time the whole-brain functional hierarchy that modulates top-down and bottom-up dependencies across spatiotemporal timescales of information processing. Clinically, this approach advances from descriptive patterns to causal, individualized mechanisms: by fitting a generative model and extracting the effective connectivity, we can identify the actual hierarchical causal drivers of non-equilibrium brain dynamics.

Future links with metabolism are suggested by ^65^, who observed that the costs required to transition within and between hierarchical levels differ significantly, with costly transitions between levels being rare and more efficient transitions within levels occurring more frequently.

Exploiting the subject-specific nature of the flow decomposition, we demonstrate that the near-critical solenoidal component carries individual signatures of temporal directionality. This aligns with the expectation that causal representations of brain function more accurately reflect underlying neural mechanisms ^17^. In this view, future steps could include exploring multivariate correlations between flow patterns and cognitive traits within each energetic range, building upon the established importance of brain–cognition relationships during resting state ^28^, which emphasized the critical dependence of broken detailed balance on the specific cognitive functions being performed.

Most importantly, our findings highlight the importance of directional, hierarchical dynamics in brain function, and our linear modeling approach provides a first step toward simultaneously characterizing the hierarchical structure emerging from non-equilibrium dynamics of multivariate neural data.

## Materials

### Data preprocessing

Our study cohort was drawn from the HCP Healthy Young Adult dataset, using the final 1200-subject release (https://www.humanconnectome.org/study/hcp-young-adult/document/1200-subjects-data-release).

For functional data, we analyzed ICA-FIX–denoised resting-state fMRI volumetric timeseries, encoded as *rfMRI_REST1_LR_hp2000_clean*, preprocessed according to the HCP functional pipeline ^67^. Preprocessing included spatial (“minimal”) preprocessing, weak high-pass temporal filtering to remove slow drifts, and ICA-FIX denoising to suppress motion- and artifact-related components.

We applied a clustered cortical parcellation comprising 62 regions (see Supplementary Methods for details), each assigned to one of the seven canonical resting-state networks defined by ^68^: visual (VIS, 5 parcels), somatosensory-motor (SOM, 6 parcels), dorsal attention (DAN, 9 parcels), salience/ventral attention (SVAN, 11 parcels), limbic (LIM, 5 parcels), cognitive control (CON, 10 parcels), and default mode (DMN, 16 parcels). Additionally, 12 subcortical and cerebellar regions (SUB) were included based on AAL2 segmentation ^69^: for each hemisphere, the thalamus proper, caudate, putamen, pallidum, cerebellum, and hippocampus. Across analyses, nodes were consistently ordered by RSN, with left-hemisphere nodes preceding right-hemisphere nodes.

To mitigate motion artifacts, we constructed a binary temporal mask for each subject, flagging volumes with framewise displacement (FD) > 0.4 mm ^70^. This mask was incorporated into sDCM estimation, down-weighting high-motion frames in the inference procedure. Time series were additionally despiked using the *icatb_despike_tc* function of the GIFT toolbox.

Temporal traces were band-pass filtered (0.008–0.1 Hz) to capture the frequency range characteristic of resting-state low-frequency fluctuations ^68,4^. This range emphasizes neuronal dynamics while minimizing contributions from hemodynamic components, which extend beyond 0.1 Hz ^71^. Given sDCM’s sensitivity to noise and the high temporal resolution of HCP data, we interpolated signals by resampling to a 1 s repetition time, yielding 864 frames per subject. These timeseries were demeaned and rescaled following the recommendations of ^72^ before serving them as algorithm inputs.

Our analysis focused on the two resting-state runs acquired with LR phase encoding. The initial sample comprised 255 participants. We excluded individuals with excessive motion (i.e., <50% of volumes with FD < 0.4 mm) and those in whom sDCM failed to converge, a common outcome under high-noise conditions. The final cohort included 144 participants (65 males, 79 females; mean age = 28.94 ± 3.55 years).

At the single-subject level, the effective connectivity matrix and associated model parameters were estimated using sDCM (see Supplementary Methods for details).

## Methods

All subsequent data analysis was conducted employing code written in MATLAB R2021b and R2022b (Mathworks, USA).

### Eigendecomposition of effective connectivity

From a dynamical systems theory perspective, the effective connectivity matrix can be conceptualized as encapsulating the internal neuronal dynamics structured by its associated eigenvectors and eigenvalues ^25^. To delve into this perspective, we performed a spectral decomposition on each individual matrix using the following spectral expansion:

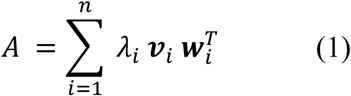

where *λ*^*i*^ denotes the *i^th^* eigenvalue of the linear system, while ***v***_*i*_ and 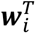 refer to the associated right and left eigenvectors, respectively. Given that we are working with an asymmetric matrix, the conversion to eigenvector coordinates may necessitate the incorporation of a complex vector space (with both real and imaginary components) ^25^. Consequently, this transformation yields non-orthogonal and complex eigenvectors, along with complex eigenvalues.

Accordingly, the temporal evolution of each neural state (i.e., the solution of the linear dynamics) can be written as

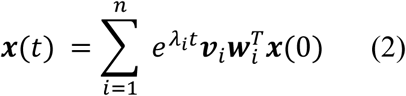

At each *λ*_*i*_, the right eigenvector ***v***_*i*_ characterizes the spatial impact of different brain regions on ***x***(*t*), while the left eigenvector 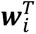 denotes the response strength to the initial state ***x***(0) ^26^. This means that the hidden neural activity can be expressed, at each time step, as a weighted (i.e., 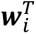***x***(0)) linear combination of the system responses (i.e., 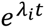) associated with the spatial pattern of each right eigenvector (i.e., ***v***_*i*_). Both Equations (1) and (2) are referred to as the dynamic modal decomposition of the dynamical system ^26^. In this view, *λ*_*i*_ and ***v***_*i*_ (real or conjugate pair when complex) describe each eigenmode providing the state coordinates to describe the spatiotemporal process of the system, whose time scales are dictated by the eigenvalues of the matrix *A*. In other words, these modes provide the low-dimensional axes along which the neural trajectories evolve at different temporal scales ^27^. The collection of functional modes available thus constitutes the functional repertoire of each EC network.

### Eigendecomposition of *S*

Adopting the thermodynamic perspective outlined by ^12^, the real and imaginary parts of each eigenvalue of the EC matrix encode two fundamental components of information flow in steady state. The real part reflects the decay rate of the dissipative flow, which manifests as a gradient flow toward higher density and counteracts the dispersive effects of random fluctuations. The imaginary part corresponds to the principal frequency of the solenoidal flow, which circulates along iso-probability contours without affecting the nonequilibrium steady-state density. According to the Helmholtz decomposition theorem, these two flows are orthogonal, and in the case of sDCM can be derived in closed form (see Supplementary Methods): the dissipative flow is mediated by the product between the factor 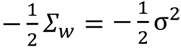 and the precision matrix *Σ*^−1^, accounting for the dispersive effect of random fluctuations, while the solenoidal flow is regulated by the product between the precision and differential cross-covariance matrix *S* (i.e., *SΣ*^−1^).

Crucially, solenoidal coupling encoded by *S* gives rise to directed macroscopic fluxes, thereby breaking detailed balance. This breaking of symmetry generates entropy and, in thermodynamic terms, establishes an irreversible “arrow of time” ^29^.

On the other hand, the link between eigenvalues and dissipative/solenoidal components also provides a direct means of quantifying the kinetic energy of the overall dynamic flow in terms of the eigenvalues of the EC matrix, disentangling how individual modes stabilize or circulate activity within the system^12^:

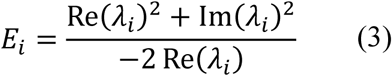

where the real (i.e., 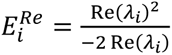) and the imaginary parts (i.e., 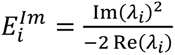) refer to the dissipative and non-dissipative energy component of each eigenstate, respectively.

Because each subject’s EC matrix yields a unique set of eigenmodes, we implemented a data-driven partitioning of the eigenspace to identify different energy ranges. For each subject, the AUC defined by 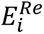 and 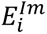 was split into two equal parts, following a procedure similar to that used for real eigenmodes in ^7^. Combining both partitions in the complex plane defined four energy-based regimes:

- R1, with 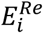 and 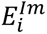 both below the AUC cutoff;
- R2, with 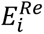 above and 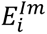 below the AUC cutoff;
- R3, with both 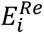 and 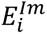 above the AUC cutoff;
- R4, with 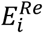 below and 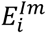 above the AUC cutoff.

_*i*_Note that for each subject we discarderd all fast eigenmodes whose real part was < -0.7: this comes from considering that, given a resolution time of 1 s, we are able to reliably estimate a half-life of roughly 1 s, which corresponds to a decay rate 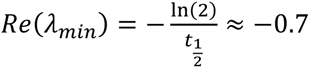

Building on this interpretation, the spectral framework introduced in the previous section allows the decomposition of *S* within the eigenmode coordinate system, enabling the quantification of each mode’s contribution to solenoidal interactions. To examine the distribution of connectivity patterns captured by *S* across the energetic ranges, we projected both matrices onto eigenvector coordinates across different timescales.

After identifying the set of eigenvalues and the associated right and left eigenvectors characterizing each energetic range *k* as 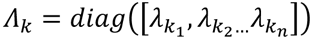, 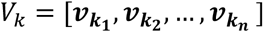 and 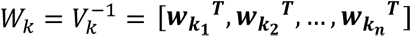 (i.e., the *k*_1_, *k*_2_, …, *k*_*n*_ rows of 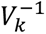) for *k=*[1,2,3, we thus performed the projection of *S* on the associated coordinates according to the following formula:

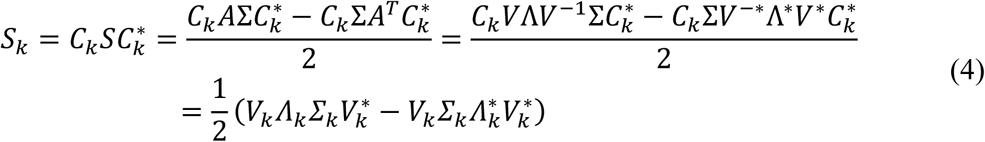

where *C*_*k*_ = *V*_*k*_*W*_*k*_ and Σ_*k*_ satifies the following Lyapunov equality

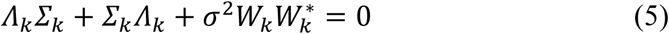

Although the notation involves complex conjugate transpose (*), eigenvalues and eigenvectors always occur in conjugate pairs, ensuring real-valued matrices. One subject was excluded at this stage because their spectral profile did not cover all energetic regimes, leaving a total of 143 subjects.

### Inference of hierarchies from *S*

To investigate temporal causality encoded in the differential cross-covariance matrices, we sought to elucidate hierarchical information processing by constructing a directed acyclic graph (DAG) for each subject. Asymmetries in the structure of *S* imply preferential information flow, thereby encoding hierarchical reorganization.

Our first step was to isolate positive entries (*S* > 0) in order to identify column-wise sender nodes. We then applied an iterative sparsification procedure: in each residual cycle, the weakest link was removed until all cycles were eliminated. This heuristic process was repeated 100 times for each subject, and only the strongest connections consistently retained across iterations contributed to the final subject-level DAG.

Each DAG was subsequently topologically sorted to assign a unique ranking of brain nodes. In cases of ambiguity, nodes were ordered by descending outdegree; ties were further resolved using outstrength. At the group level, hierarchical rankings were derived by averaging the ranking distributions across subjects for each ROI and ordering them in ascending order from source to sink (see violin plots in Supplementary Fig. S3 and S13). To assess the role of different energetic ranges in shaping cortical hierarchy, we tested the significance of the network rankings using the Jonckheere–Terpstra trend test (α=0.01).

To statistically evaluate the structure of the inferred DAGs, we compared the number and cumulative weight of links removed during cycle-breaking with those obtained from a null model. The null model was constructed by generating subject-level randomized solenoidal matrices (*S*^*rand*^) that preserved the antisymmetric structure of the empirical *S* matrices. Specifically, for each node pair (*i,j*), if either S_*ij*_ or S_*ji*_was nonzero, the interaction direction was reassigned randomly according to a uniform distribution, while ensuring (*S*^*rand*^ = -*S*^*rand*^). Each randomized matrix was then subjected to the same sparsification procedure as the empirical data, enabling statistical comparison of the number and weight of removed links between real and null networks (Wilcoxon rank-sum test, α=0.05).

### Fingerprinting analysis of *S*

To assess the fingerprinting capacity of *S* across energetic ranges, we adapted a methodology commonly employed in functional connectivity fingerprinting ^35^, based on the construction of an identifiability matrix. This similarity matrix, defined as the Pearson correlation between each pair of connectivity profiles from test and retest sessions, captures both self-similarity (main diagonal) and similarity between different subjects (off-diagonal).

We quantified differential identifiability (*I*_*diff*_), under the assumption that connectivity profiles should be more similar within the same subject across sessions than between different subjects:

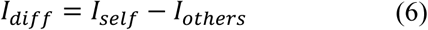

where *I*_*self*_ =< *I*_*ii*_ > represent the average of the main diagonal elements of the identifiability matrix *I* (i.e., correlation values between visits of same subjects), while *I*_*others*_ =< *I*_*ij*_ > defines the average of the off-diagonal elements of *I* (i.e., correlation between visits of different subjects). Finally, we measured the success rate (SR) of the identification procedure as the percentage of cases with higher within-(*I*_*self*_) vs. between-subjects (*I*_*others*_) test–retest reliability ^35^. Differences in *I*_*diff*_ and SR across ranges were tested using repeated-measures ANOVA with subject as a within-subject factor, followed by Tukey–Kramer post hoc correction for multiple comparisons (*α*=0.05).

Statistical significance of both differential identifiability and success rate mean values was evaluated using permutation testing ^36^. At each iteration, subjects’ test–retest connectomes were randomly shuffled, and the corresponding identifiability metrics were recomputed. This procedure was repeated 1,000 times to obtain a nonparametric null distribution of average *I*_*diff*_ and SR. Observed values were then compared against this distribution, and p-values were determined as the proportion of permutations exceeding the empirical scores (99^th^ percentile threshold).

Beyond whole-brain measures, we also assessed edgewise subject identifiability by computing the intraclass correlation coefficient (ICC) ^73^:

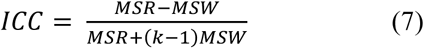

where (MSR) is the mean square between subjects, (MSW) is the mean square of residual variance, and (k=2) is the number of sessions. A high ICC indicates that test–retest reliability for a given edge is subject-specific, whereas a low ICC suggests variability dominated by session-to-session fluctuations or noise.

From the edgewise ICC values, we constructed a symmetric ICC matrix of size N=74. We further explored network-level fingerprinting by averaging ICC values across intra- and inter-network connections, yielding (8×8) ICC matrices corresponding to the cortical and subcortical network parcellation. In this framework, higher ICC reflects greater fingerprinting strength at the network level.

Finally, nodal fingerprinting strength was quantified by summing column-wise values of the ICC matrix, providing a region-wise measure of fingerprinting capacity. Distributions of nodal strength across energetic ranges were visualized using brain renderings. To enhance interpretability, values were thresholded at the 1^st^–99^th^ percentiles, following the procedure described in ^36^.

### Statistical analyses

All statistical analyses were conducted in MATLAB R2021b/R2022b. Unless otherwise stated, tests were two-tailed with α = 0.05. Replicates are defined as independent subject-level estimates of effective connectivity (EC), solenoidal flow (S), eigenmodes, or derived graph metrics. Exact sample sizes were: n = 143 for EC/S-based analyses and n = 142 for structure–function (fiber length) analyses (one subject excluded due to tractography failure). For representative matrices or eigenmodes, the number of times results were reproduced across subjects is reported in the corresponding figure legends.

Effective connectivity was estimated using sparse Dynamic Causal Modeling (sDCM) with automated sparsity-inducing regularization and motion-weighted likelihood estimation. Model accuracy was evaluated using normalized root mean squared error (NRMSE) and correlations between empirical and simulated functional connectivity. All analyses used a consensus-clustered Schaefer 100-area cortical atlas to ensure spatial homogeneity and computational reproducibility. Full implementation details are provided in the Methods and Supplementary Methods for independent replication.

For hierarchy estimation, directed acyclic graphs (DAGs) were derived from S matrices using iterative sparsification (100 repetitions per subject). Trends across energetic regimes were tested using the Jonckheere–Terpstra test (one-tailed). Comparisons with null-model S matrices used the Wilcoxon rank-sum test (two-tailed).

Fingerprinting analyses used test–retest S matrices (n = 143, k = 2 sessions). Differential identifiability (*I_diff_*) and success rate (SR) were compared across energetic regimes using repeated-measures ANOVA; F values, degrees of freedom, and p-values are reported in the Results. Post hoc comparisons were corrected using the Tukey–Kramer test. Statistical significance of *I_diff_* and SR relative to chance was computed using 1,000-permutation testing (one-tailed). Edgewise reliability was measured using the intraclass correlation coefficient (ICC; k = 2 sessions), and network- and nodal-level ICC scores were summarized descriptively.

For structure–function relationships, DAG connections were classified as short-, middle-, or long-range using group-level fiber-length quartiles. Differences in proportions across energetic regimes were assessed using the Friedman test (two-tailed; α = 0.01), followed by corrected post hoc comparisons.

## Data availability

The data that support the findings of this study are available in the Human Connectome Project platform (db.humanconnectome.org). Identifiers of the included subjects are reported here: [https://github.com/FairUnipd/Effective_connectivity_eigenmodes/blob/main/data/DATA_EC_HCP.mat].

## Code availability

Example code implementing these analyses described in the study is available at: https://github.com/FairUnipd/Effective_connectivity_eigenmodes.git. The computation of *I_diff_*, SR and ICC across ranges was facilitated by adapting resources from https://github.com/eamico/Clinical_fingerprinting ^60^ and https://github.com/ss1913/fingerprints_alzheimer ^74^. The code to apply the Jonckheere–Terpstra test was adapted from https://github.com/dnafinder/jttrend.

## Author contributions

Giorgia Baron, Giacomo Baggio and Danilo Benozzo designed the research. Massimiliano Facca performed the extraction of structural connectomes. Giorgia Baron analysed the data. Giorgia Baron, Giacomo Baggio, Massimiliano Facca, Danilo Benozzo, Sandro Zampieri and Alessandra Bertoldo interpreted the results. Alessandro Chiuso provided the sparse DCM model. Giorgia Baron wrote the manuscript. All authors revised the manuscript.

## References

1. Vohryzek, J., Sanz-Perl, Y., Kringelbach, M. L. & Deco, G. Human brain dynamics are shaped by rare long-range connections over and above cortical geometry. Proc. Natl. Acad. Sci. 122, e2415102122 (2025).

2. Geli, S. M., Lynn, C. W., Kringelbach, M. L., Deco, G. & Sanz Perl, Y. Non-equilibrium whole-brain dynamics arise from pairwise interactions. Cell Rep. Phys. Sci. 6, 102464 (2025).

3. Glomb, K. et al. Functional harmonics reveal multi-dimensional basis functions underlying cortical organization. Cell Rep. 36, 109554 (2021).

4. Margulies, D. S. et al. Situating the default-mode network along a principal gradient of macroscale cortical organization. Proc. Natl. Acad. Sci. 113, 12574–12579 (2016).

5. Atasoy, S., Donnelly, I. & Pearson, J. Human brain networks function in connectome-specific harmonic waves. Nat. Commun. 7, 10340 (2016).

6. Luppi, A. I. et al. Distributed harmonic patterns of structure-function dependence orchestrate human consciousness. *Commun*. Biol. 6, 117 (2023).

7. Preti, M. G. & Van De Ville, D. Decoupling of brain function from structure reveals regional behavioral specialization in humans. Nat. Commun. 10, 4747 (2019).

8. Wang, M. B., Owen, J. P., Mukherjee, P. & Raj, A. Brain network eigenmodes provide a robust and compact representation of the structural connectome in health and disease. PLOS Comput. Biol. 13, e1005550 (2017).

9. Nartallo-Kaluarachchi, R., Kringelbach, M. L., Deco, G., Lambiotte, R. & Goriely, A. Nonequilibrium physics of brain dynamics. Preprint at 10.48550/arXiv.2504.12188 (2025).

10. Kringelbach, M. L., Sanz Perl, Y. & Deco, G. The Thermodynamics of Mind. Trends Cogn. Sci. 28, 568–581 (2024).

11. Van Essen, D. C. et al. The Human Connectome Project: a data acquisition perspective. NeuroImage 62, 2222–2231 (2012).

12. Friston, K. J. et al. Parcels and particles: Markov blankets in the brain. Netw. Neurosci. 5, 211–251 (2021).

13. Deco, G. et al. Rare long-range cortical connections enhance human information processing. Curr. Biol. CB 31, 4436–4448.e5 (2021).

14. Hillebrand, A. et al. Direction of information flow in large-scale resting-state networks is frequency-dependent. Proc. Natl. Acad. Sci. 113, 3867–3872 (2016).

15. Atasoy, S., Deco, G., Kringelbach, M. L. & Pearson, J. Harmonic Brain Modes: A Unifying Framework for Linking Space and Time in Brain Dynamics. Neurosci. Rev. J. Bringing Neurobiol. Neurol. Psychiatry 24, 277–293 (2018).

16. Alamia, A. & VanRullen, R. Alpha oscillations and traveling waves: Signatures of predictive coding? PLOS Biol. 17, e3000487 (2019).

17. Bolton, T. A. W., Van De Ville, D., Amico, E., Preti, M. G. & Liégeois, R. The arrow-of-time in neuroimaging time series identifies causal triggers of brain function. Hum. Brain Mapp. 44, 4077–4087 (2023).

18. Camassa, A. et al. The temporal asymmetry of cortical dynamics as a signature of brain states. Sci. Rep. 14, 24271 (2024).

19. Fousek, J. et al. Symmetry breaking organizes the brain’s resting state manifold. Sci. Rep. 14, 31970 (2024).

20. Shinozuka, K. et al. LSD flattens the hierarchy of directed information flow in fast whole-brain dynamics. Imaging Neurosci. 3, imag_a_00420 (2025).

21. Pines, A. et al. Development of top-down cortical propagations in youth. Neuron 111, 1316–1330.e5 (2023).

22. Finn, E. S. et al. Functional connectome fingerprinting: identifying individuals using patterns of brain connectivity. Nat. Neurosci. 18, 1664–1671 (2015).

23. Van De Ville, D., Farouj, Y., Preti, M. G., Liégeois, R. & Amico, E. When makes you unique: Temporality of the human brain fingerprint. Sci. Adv. 7, eabj0751 (2021).

24. Griffa, A., Amico, E., Liégeois, R., Van De Ville, D. & Preti, M. G. Brain structure-function coupling provides signatures for task decoding and individual fingerprinting. NeuroImage 250, 118970 (2022).

25. Lambiotte, R. & Schaub, M. T. Modularity and Dynamics on Complex Networks. (Cambridge University Press, Cambridge, 2022). doi:10.1017/9781108774116.

26. Griffith, T. D. & Hubbard, J. E. System identification methods for dynamic models of brain activity. Biomed. Signal Process. Control 68, 102765 (2021).

27. McIntosh, A. R. & Jirsa, V. K. The hidden repertoire of brain dynamics and dysfunction. Netw. Neurosci. 3, 994–1008 (2019).

28. Lynn, C. W., Cornblath, E. J., Papadopoulos, L., Bertolero, M. A. & Bassett, D. S. Broken detailed balance and entropy production in the human brain. Proc. Natl. Acad. Sci. 118, e2109889118 (2021).

29. Deco, G. et al. The arrow of time of brain signals in cognition: Potential intriguing role of parts of the default mode network. Netw. Neurosci. 7, 966–998 (2023).

30. Casti, U., et al. Dynamic Brain Networks with Prescribed Functional Connectivity. Preprint at 10.48550/arXiv.2310.07262 (2023).

31. Benozzo, D. et al. Analyzing asymmetry in brain hierarchies with a linear state-space model of resting-state fMRI data. Netw. Neurosci. 8, 965–988 (2024).

32. Lin, T. W., Das, A., Krishnan, G. P., Bazhenov, M. & Sejnowski, T. J. Differential Covariance: A New Class of Methods to Estimate Sparse Connectivity from Neural Recordings. Neural Comput. 29, 2581–2632 (2017).

33. Luo, J. & Magee, C. L. Detecting evolving patterns of self-organizing networks by flow hierarchy measurement. Complexity 16, 53–61 (2011).

34. Deco, G., Sanz Perl, Y. & Kringelbach, M. L. Complex harmonics reveal low-dimensional manifolds of critical brain dynamics. *Phys*. Rev. E 111, 014410 (2025).

35. Amico, E. & Goñi, J. The quest for identifiability in human functional connectomes. Sci. Rep. 8, (2018).

36. Sareen, E. et al. Exploring MEG brain fingerprints: Evaluation, pitfalls, and interpretations. NeuroImage 240, 118331 (2021).

37. Medaglia, J. D. et al. Functional Alignment with Anatomical Networks is Associated with Cognitive Flexibility. *Nat*. Hum. Behav. 2, 156–164 (2018).

38. Christodoulou, G. & Vogels, T. The Eigenvalue Value (in Neuroscience). Preprint at 10.31219/osf.io/evqhy (2022).

39. Friston, K. J., Kahan, J., Razi, A., Stephan, K. E. & Sporns, O. On nodes and modes in resting state fMRI. NeuroImage 99, 533–547 (2014).

40. Bolt, T. et al. A parsimonious description of global functional brain organization in three spatiotemporal patterns. Nat. Neurosci. 25, 1093–1103 (2022).

41. Bohon, M. D., Orchini, A., Bluemner, R., Paschereit, C. O. & Gutmark, E. J. Dynamic mode decomposition analysis of rotating detonation waves. Shock Waves 31, 637–649 (2021).

42. Zhang, J. et al. Intrinsic Functional Connectivity is Organized as Three Interdependent Gradients. Sci. Rep. 9, 15976 (2019).

43. Feldman, H. & Friston, K. Attention, Uncertainty, and Free-Energy. Front. Hum. Neurosci. 4, (2010).

44. Liu, Z.-Q. et al. Time-resolved structure-function coupling in brain networks. *Commun*. Biol. 5, 532 (2022).

45. Gu, Y. et al. Brain Activity Fluctuations Propagate as Waves Traversing the Cortical Hierarchy. Cereb. Cortex N. Y. N 1991 31, 3986–4005 (2021).

46. Mitra, A., et al. Human cortical–hippocampal dialogue in wake and slow-wave sleep. Proc. Natl. Acad. Sci. 113, E6868–E6876 (2016).

47. Stanovich, K. E. & West, R. F. Individual differences in reasoning: implications for the rationality debate? Behav. Brain Sci. 23, 645–665; discussion 665-726 (2000).

48. Sadaghiani, S., Brookes, M. J. & Baillet, S. Connectomics of human electrophysiology. NeuroImage 247, 118788 (2022).

49. Lombardi, F., Pepić, S., Shriki, O., Tkačik, G. & De Martino, D. Statistical modeling of adaptive neural networks explains co-existence of avalanches and oscillations in resting human brain. Nat. Comput. Sci. 3, 254–263 (2023).

50. Rosanova, M. et al. Natural frequencies of human corticothalamic circuits. J. Neurosci. Off. J. Soc. Neurosci. 29, 7679–7685 (2009).

51. Mahjoory, K., Schoffelen, J.-M., Keitel, A. & Gross, J. The frequency gradient of human resting-state brain oscillations follows cortical hierarchies. eLife 9, e53715 (2020).

52. Koller, D. P., Schirner, M. & Ritter, P. Human connectome topology directs cortical traveling waves and shapes frequency gradients. Nat. Commun. 15, 3570 (2024).

53. Yang, Y. et al. Enhanced brain structure-function tethering in transmodal cortex revealed by high-frequency eigenmodes. Nat. Commun. 14, 6744 (2023).

54. Pang, J. C. et al. Geometric constraints on human brain function. Nature 618, 566–574 (2023).

55. Luppi, A. I. et al. Competitive interactions shape brain dynamics and computation across species. BioRxiv Prepr. Serv. Biol. 2024.10.19.619194 (2024) doi:10.1101/2024.10.19.619194.

56. Jo, Y., Faskowitz, J., Esfahlani, F. Z., Sporns, O. & Betzel, R. F. Subject identification using edge-centric functional connectivity. NeuroImage 238, 118204 (2021).

57. Cassone, B. et al. TR(Acking) Individuals Down: Exploring the Effect of Temporal Resolution in Resting-State Functional MRI Fingerprinting. Hum. Brain Mapp. 46, e70125 (2025).

58. Sorrentino, P. et al. Brain fingerprint is based on the aperiodic, scale-free, neuronal activity. NeuroImage 277, 120260 (2023).

59. Alderson, T. H., Bokde, A. L. W., Kelso, J. A. S., Maguire, L. & Coyle, D. Metastable neural dynamics underlies cognitive performance across multiple behavioural paradigms. Hum. Brain Mapp. 41, 3212–3234 (2020).

60. Sorrentino, P. et al. Clinical connectome fingerprints of cognitive decline. NeuroImage 238, 118253 (2021).

61. Nakuci, J. et al. Within-subject reproducibility varies in multi-modal, longitudinal brain networks. Sci. Rep. 13, 6699 (2023).

62. Noble, S. et al. Influences on the Test–Retest Reliability of Functional Connectivity MRI and its Relationship with Behavioral Utility. Cereb. Cortex 27, 5415–5429 (2017).

63. Younger, D. Minimum Feedback Arc Sets for a Directed Graph. IEEE Trans. Circuit Theory 10, 238–245 (1963).

64. Nozari, E. et al. Macroscopic resting-state brain dynamics are best described by linear models. *Nat*. Biomed. Eng. 8, 68–84 (2024).

65. Ceballos, E. G. et al. The control costs of human brain dynamics. Netw. Neurosci. 9, 77–99 (2025).

66. Muldoon, S. F. et al. Stimulation-Based Control of Dynamic Brain Networks. PLOS Comput. Biol. 12, e1005076 (2016).

67. Glasser, M. F. et al. The minimal preprocessing pipelines for the Human Connectome Project. NeuroImage 80, 105–124 (2013).

68. Yeo, B. T. T. et al. The organization of the human cerebral cortex estimated by intrinsic functional connectivity. J. Neurophysiol. 106, 1125–1165 (2011).

69. Rolls, E. T., Joliot, M. & Tzourio-Mazoyer, N. Implementation of a new parcellation of the orbitofrontal cortex in the automated anatomical labeling atlas. NeuroImage 122, 1–5 (2015).

70. Power, J. D. et al. Methods to detect, characterize, and remove motion artifact in resting state fMRI. NeuroImage 84, 320–341 (2014).

71. Tong, Y., Hocke, L. M. & Frederick, B. B. Low Frequency Systemic Hemodynamic “Noise” in Resting State BOLD fMRI: Characteristics, Causes, Implications, Mitigation Strategies, and Applications. Front. Neurosci. 13, (2019).

72. Zeidman, P. et al. A guide to group effective connectivity analysis, part 1: First level analysis with DCM for fMRI. NeuroImage 200, 174–190 (2019).

73. McGraw, K. O. & Wong, S. P. Forming inferences about some intraclass correlation coefficients. Psychol. Methods 1, 30–46 (1996).

74. Stampacchia, S. et al. Fingerprints of brain disease: connectome identifiability in Alzheimer’s disease. *Commun*. Biol. 7, 1169 (2024).

