## Supplementary material for "Eigenmode decomposition of asymmetries in whole-brain effective connectivity reveals multiscale hierarchical dynamics"

### Supplementary Information for: Eigenmode decomposition of asymmetries in whole-brain effective connectivity reveals multiscale hierarchical dynamics

**Authors:** Giorgia Baron[1,2], Giacomo Baggio[1], Massimiliano Facca[1], Danilo Benozzo[1,3], Alessandro Chiuso[1], Sandro Zampieri[1], Alessandra Bertoldo[1,4*]

[1]Department of Information Engineering, University of Padova, Padova, Italy
[2] Defitech Chair of Clinical Neuroengineering, Neuro-X Institute (INX), École Polytechnique Fédérale de Lausanne (EPFL), 1202 Geneva, Switzerland; Defitech Chair of Clinical Neuroengineering, INX, EPFL Valais, Clinique Romande de Réadaptation, 1950 Sion, Switzerland
[3] Department of Brain and Behavioral Sciences, University of Pavia, Italy
[4]Padova Neuroscience Center, University of Padova, Padova, Italy

**Correspondence and requests for materials should be addressed to Alessandra Bertoldo**
Department of Information Engineering, University of Padova, Padova, Italy
Padova Neuroscience Center, University of Padova, Padova, Italy

https://orcid.org/0000-0002-6262-6354

### Supplementary Methods

#### Data collection

Our study cohort is derived from the HCP Healthy Young Adult dataset, available in its final release at https://www.humanconnectome.org/study/hcp-young-adult/document/1200-subjects-data-release.

This extensive public dataset comprises individuals aged between 22 and 35 years, all without any psychiatric or neurological disorders, as documented by [1]. The acquisition parameters and minimal preprocessing details for these data are outlined in [2] and involve high spatial (2 mm isotropic) and temporal (TR = 0.72 s) resolution multi-band fMRI. The HCP subjects were scanned on a specialized Siemens 3T "Connectome Skyra" located at Washington University in St. Louis, using a standard 32-channel Siemens receive head coil.

Each participant underwent a comprehensive set of MRI scans, including T1w and T2w structural imaging and nearly 2 hours of resting-state and task fMRI. Specifically, for the resting-state sessions, rfMRI data were acquired in four runs of approximately 15 minutes each, split between two sessions on two different days. Participants maintained an open-eyed, relaxed fixation state during data acquisition. The sequence parameters for each run were as follows: gradient-echo EPI, TR = 720 ms, TE = 33.1 ms, flip angle = 52°, FOV = 208 × 180 mm, voxel size = 2 mm isotropic, frames = 1200. Within each session, oblique axial acquisitions alternated between phase encoding in a right-to-left (RL) direction in one run and phase encoding in a left-to-right (LR) direction in the other run.

#### Cortical atlas definition

The cortical parcellation employed in this study comprises 62 regions derived from the Schaefer 100-area atlas [3] organized into 7 [4] functional networks.

The decision to cluster the original atlas stems from the necessity to maintain the computational load of sparse DCM at a manageable level [5]. The clustered atlas was created by merging areas that were already functionally grouped in the atlas, as detailed by [6], while keeping other areas separate. Specifically, a consensus clustering procedure, similar to that described by [7], was employed to reduce the number of cortical parcels derived from the Schaefer atlas. This framework employs both base and consensus clustering methods to achieve robust and stable clusters. Initially, various groupings are generated for each individual dataset using a base clustering method. Then, consensus clustering is applied to these groupings to produce stable clusters across individuals. To account for hemodynamic differences across spatially distant parcels, this procedure is selectively performed on subsets of adjacent cortical regions within the same functional network. This additional constraint ensures that only functionally homogeneous and spatially contiguous parcels are grouped together.

For sets composed of only two contiguous ROIs, the objective criteria for determining the optimal number of clusters cannot be applied. In these cases, grouping is performed if the pair Pearson's correlation is, on average, greater than the mean correlation within other clusters. This approach

ensures the hemodynamic consistency of each cluster, reflecting the whole-brain vascular complexity [8,9].
The Nifti file corresponding to the obtained functional parcellation and the associated table are available at this link: https://github.com/FairUnipd/Effective_connectivity_eigenmodes.git.

**Sparse DCM**

For each subject, the resting-state hemodynamic responses and the EC matrix were inferred using the sparse DCM (sDCM) framework [5].
The original DCM framework [10] operates as a generative state-space model. In this model, the neural state *x(t)*, representing the activity of brain nodes (i.e., whole-brain regions), follows a coupled system of ordinary differential equations that describe the interactions among neural components. Simultaneously, the output model maps neuronal activity to the measured BOLD signal *y(t)* through the hemodynamic response function (HRF):

$$\frac{d\boldsymbol{x}}{dt} = A\boldsymbol{x}(t) + \boldsymbol{\nu}(t) \tag{1}$$

$$\boldsymbol{y}(t) = h(\boldsymbol{x}(t); \theta_h) + \boldsymbol{e}(t) \tag{2}$$

where $\boldsymbol{x}(t) = [x_1(t), x_2(t), \dots x_n(t)]^T$ is the hidden neural activity characterizing the configuration in the state space of the brain regions at time *t*, *A* represents the effective connectivity matrix, $\boldsymbol{y}(t)$ is the BOLD fMRI response at time *t* and $\theta_h$ denotes collectively a set of biophysical parameters regulating the hemodynamic response (which is modelled with the Balloon-Windkessel model). Moreover, the variable $\boldsymbol{\nu}(t)$, expressing the stochastic intrinsic brain fluctuations, and $\boldsymbol{e}(t)$, denoting the observation noise, are assumed to be Gaussian variables with zero mean and diagonal covariance matrices $\sigma^2 I_n$ ($I_n$ = the identity matrix of size *n*) and $R=diag(\lambda_1, \lambda_2, \dots \lambda_n)$, respectively.
In contrast to traditional DCM approaches that assume a fixed HRF, sDCM introduces a statistical linearization and discretization procedure of Eq. 1 and 2. This approach is motivated by the relatively low temporal resolution of fMRI data, typically ranging from 0.5 to 3 seconds, and the assumption that the hemodynamic response can be modeled as a finite impulse response, where the neuronal state serves as the input and the BOLD signal as the output [5].
Concerning HRF, this procedure transforms empirical priors on the physiological parameters of the nonlinear hemodynamic response model [11] into a hemodynamic prior, which is then used in the HRF estimation process. This approach provides a whole-brain characterization of the spatial pattern of hemodynamic variability, as demonstrated in our prior work [6].
Regarding connectivity parameters, this variant of DCM incorporates a sparsity-inducing mechanism that automatically prunes irrelevant connections, eliminating the need to manually select between competing network structures. The model inversion and parameter optimization are performed through an expectation-maximization (EM) algorithm combined with an iterative reweighted procedure, as detailed in [5].
As briefly mentioned in the Methods section, to improve the robustness of the model, the algorithm has been enhanced to accommodate the signal reliability of fMRI temporal frames, which are often

affected by motion artifacts. This is achieved by incorporating a binary temporal mask as a weighting measure during the estimation process. The model also considers that the measurement noise (denoted as $e$ in Equation (12) in [5]) follows a Gaussian white noise distribution with a stationary diagonal covariance matrix $R$. However, this assumption could lead to potential overfitting when the input signal undergoes band-pass filtering, which removes higher-frequency components, possibly leading to quasi-zero estimates of noise variance for each brain region. To address this limitation, we introduced a heuristic grid search, exploring a range of values for the noise variance across reasonable fractions of the sample variance of $y$ (e.g., from 1/2 to 1/100). The optimal value was selected based on two metrics: Pearson's correlation between the triangular components of the empirical functional connectivity (eFC) and the simulated functional connectivity (sFC), and the similarity between the distributions of their respective dynamic versions, as outlined in [12]. The noise variance was set to 1/10 for both runs.

For more detailed insights into model implementation and the inference procedure, readers are referred to [5]. Using the sDCM framework, we estimated the EC matrix and 74 ROI-specific HRF profiles (representing 62 cortical and 12 subcortical/cerebellar nodes) at the individual level, providing a comprehensive characterization of spatial variations in neurovascular coupling.

Then, to evaluate the model fit, we applied two summary metrics to assess the reliability of the inferred state-space DCM models and their corresponding parameters. First, for each subject and ROI, we calculated the normalized root mean squared error (NRMSE) between the actual BOLD signal and the signal predicted by the individual-level model. Normalization of the RMSE allows for comparisons across datasets or models with varying scales. We used interquartile range normalization (i.e., the difference between the 25$^{th}$ and 75$^{th}$ percentiles) to reduce sensitivity to outliers. Second, we examined the distribution of Pearson correlation coefficients between eFC and sFC across subjects. Our models achieved consistently good fits (e.g., Run1: whole-brain median NRMSE = 17.78%, median eFC–sFC correlation =0.95; Run2: median NRMSE = 17.97%, median eFC–sFC correlation =0.94), suggesting that sDCM effectively captures the physiological variability and dynamic structure underlying the observed BOLD signals.

#### Extraction of *S* from EC

As discussed in [13], the kinetic energy of functional information flow can be decomposed into dissipative and non-dissipative components, corresponding to the emergence of the curl-free dissipative and divergence-free solenoidal contributions, respectively. The dissipative component manifests as a gradient flow, directing towards increasing density and counteracting the dispersive impact of random fluctuations. In contrast, the solenoidal component circulates along iso-probability contours, exerting no influence on the nonequilibrium steady-state density.

As demonstrated in a recent work [14], these two energetic flow components can be statistically derived from EC in a closed-form. More specifically, when the Hurwitz stability of $A$ (i.e., the EC matrix) is granted, it may be decomposed as follows [15]:

$$A = \left(-\frac{1}{2}\Sigma_w + S\right)\Sigma^{-1} \tag{3}$$

On one hand, the matrix $\Sigma_w$ (i.e., the differential auto-covariance) corresponds to the positive definite stationary form of the matrix $Q$ defined in [5], namely, $\Sigma_w = E[\boldsymbol{w}(t)\boldsymbol{w}(t)^\top]$. Since we assumed uncorrelated noise across nodes, $\Sigma_w = \sigma^2 I_n$ where $\sigma^2$ represents the variance of each brain endogenous fluctuations and $I_n$ is the identity matrix of size $n$. On the other hand, the inverse of the covariance matrix $\Sigma^{-1}$ (also known as precision matrix) derives from inverting the steady-state covariance matrix $\Sigma$ defined as:

$$\Sigma \coloneqq \lim_{t\to\infty} E\,[\boldsymbol{x}(t)\boldsymbol{x}(t)^\top] \tag{4}$$

for which the algebraic Lyapunov equation holds [15]:

$$A\Sigma + \Sigma A^\top + \Sigma_w = 0 \tag{5}$$

As extensively discussed in [16], $-\Sigma^{-1}$ is related to the pairwise partial covariance between each pair of neural nodes, since it captures only "direct" statistical dependencies by discarding the effects of mediators. From a thermodynamic perspective, the precision matrix also reflects how sharply the probability density is concentrated around certain states, effectively shaping the curvature of the underlying potential energy landscape on which the gradient descent takes place to counteract the dispersive effect of noise [17].

While $\Sigma_w$ and $\Sigma^{-1}$ are both symmetric, the asymmetry of $A$ relies on the nature of the skew-symmetric matrix $S = -S^T$, thus introducing a solenoidal regime in the energy landscape of brain functional dynamics associated with the non-dissipative component of the functional flow's kinetic energy. Since Eq. 3 mathematically parametrizes all EC matrices that give the same $\Sigma$ for any matrix $S$, this defines a one-to-one mapping between $A$ and $(\Sigma, S)$ given $\Sigma_w$, thus leading to compute $S$ as:

$$S \coloneqq \frac{1}{2}(A\Sigma - \Sigma A^\top) \tag{6}$$

In statistical terms, $S$ can be referred to as the differential cross-covariance matrix [18]. Under the assumption of a diagonal noise covariance structure, it follows that any out-diagonal element $(i, j)$ of $S$ corresponds to the differential covariance $\lim_{t\to\infty} E\left[\dot{x}_i(t)x_j(t)^\top\right] = -\frac{1}{2}\Sigma_w + S$, which is antisymmetric (or skew-symmetric) by definition (see Lemma 3, Appendix A in [18]). It follows that the time-lag covariance matrix $S$ provides a direct interpretation of which node behaves as a source and which as a sink. Specifically, if $S_{ij} > 0$ (and $S_{ji} < 0$ for the skew-symmetry of $S$), the node j is considered as source, meaning that information is flowing from node j to node i. As a result, this statistical interpretation provides a straightforward method to ascertain the sender or receiver nature of each neural node. By summing the values of matrix $S$ column-wise, we can quantify the extent to which each node predominantly functions as a source (positive sum) or a sink (negative sum).

Specifically, when $S$ equals zero, the eigenvalues of $A$ are real, due to the symmetric nature of the resulting EC matrix. Conversely, as the entries of $S$ increase, the eigenvalues of A transition to a complex state, underscoring the profound impact of S on the system's non equilibrium.

**Comparative analysis of $S$ and the dominant eigenvector**

Given the tight relationship between EC flow patterns and their associated eigenspaces, we sought to clarify how the spatial configuration of eigenvectors contributes to the emergence of network-level dynamics across distinct energetic regimes. This analysis aimed to provide a more mechanistic and physically interpretable account of variability in the complex eigenvector structure, particularly the wave-like propagation patterns encoded in the spatiotemporal structure of *S*.

To formalize this intuition, we further developed the expression of the system's temporal evolution along the $i^{th}$ eigenmode. Specifically, each region's response projected on $\boldsymbol{v}_i$ can be described in real terms by decomposing the complex eigenvector $\boldsymbol{v}_i$ into its modulus and phase components:

$x_{ROI}(t) = 2\rho_{ROI}e^{\alpha t}(A_1\cos(\beta t + \varphi_{ROI}) - A_2\sin(\beta t + \varphi_{ROI}))$ , where $A_1$ and $A_2$ are real constants dependent on the product $\boldsymbol{w}_i^T\boldsymbol{x}(0)$, $\alpha = \mathrm{Re}(\lambda_i)$, $\beta = \mathrm{Im}(\lambda_i)$, while $\rho_{ROI}$ and $\varphi_{ROI}$ represent the ROI-level modulus and phase of $\boldsymbol{v}_i$. This formulation clearly indicates that, on one hand, the magnitude $\rho_{ROI}$ governs the amplitude of neural fluctuations, identifying subpopulations of ROIs exhibiting consistent behavior at different scales of amplitude synchronization, irrespective of whether $\mathrm{Im}(\lambda_i)$=0 or $\mathrm{Im}(\lambda_i) > 0$.

On the other hand, disparities in phase $\varphi_{ROI}$ clearly delineate the temporal precedence or succession between each pair of brain regions, naturally establishing an associated arrow of time that emanates from the group of senders and points towards the group of receivers. To assess how these eigenvector-derived quantities relate to causal brain dynamics, we focused on the dominant eigenmode for each energetic range within each subject, defined as the eigenvector corresponding to the least negative real part of the eigenvalue (i.e., the slowest decaying mode). For each pair of eigenvector complex entries, we then computed all pairwise phase differences and products of modulus values between regions.

To ensure consistency and comparability across subjects and ranges, phase differences were wrapped into the interval $[-\pi, \pi]$, allowing us to systematically examine how phase lags and amplitude interactions relate to the empirically estimated directionality of information flow.

These measures were compared to the corresponding entries in the upper triangular portion of matrix *S*, capturing directed influences between regions (see Fig. S1A). Importantly, since *S* is statistically equivalent to the differential cross-covariance between time series — and thus reflects the temporal asymmetry of interactions — it offers a principled measure of directionality. This justifies our use of phase differences in interpreting *S*: when the phase difference between regions *k* and *j* ($\phi_k - \phi_j$) is negative, meaning that region *k* precedes *j* (either due to a phase lead >180° or a lag <0°), we expect $S_{kj} < 0$ to be more probable than $S_{kj} > 0$. Conversely, when the phase difference is positive, indicating that *j* precedes *k*, a positive $S_{kj}$ becomes more likely. This directional bias provides a theoretical rationale for the observed correlation between phase disparities and the sign and magnitude of entries in *S*. Moreover, as *S* represents a covariance measure, we expect that, for each entry, a relationship also exists with the product of each pair of magnitudes.

**Assessment of the fiber length underlying hierarchical structure**

To relate the inferred directed acyclic graphs (DAGs) to underlying structural connections, we analyzed the distribution of fiber lengths associated with the directed connections underlying individual DAGs. For this purpose, the HCP dataset provides high-resolution Diffusion Weighted Imaging (DWI) data (spin-echo planar imaging sequence; voxel size: 1.25 mm isotropic; TR: 5520 ms; TE: 89.5 ms; maximum b-value: 3000 s/mm²; 270 non-collinear diffusion directions; 18 B0 acquisitions). The data are provided after minimal preprocessing, including B0 intensity normalization and correction for susceptibility distortions, eddy currents, motion, and gradient nonlinearities, as well as registration to the high-resolution T1-weighted image (as described in [2]. We performed probabilistic tractography in the space of the minimally preprocessed data. Fiber orientation distributions (FODs) were estimated using Multi-shell Multi-tissue Constrained Spherical Deconvolution [19]. The high-resolution T1-weighted image was segmented using a Hybrid Surface-Volume Segmentation approach [20] to enable Anatomically-Constrained Tractography (ACT;[21]) during global fiber tracking. For each subject, we generated a whole-brain tractogram using the iFOD2 algorithm, with streamlines dynamically seeded within the white matter [22], until reaching 10 million valid streamlines. To improve the biological plausibility of the tractograms and mitigate biases inherent to probabilistic tractography, we applied Spherical-deconvolution Informed Filtering of Tractograms (SIFT2; [22]). Finally, we generated weighted and symmetric structural connectivity matrices by summing the SIFT2 streamline weights, as well as corresponding weighted and symmetric fiber-length matrices.

To minimize the influence of unreliable connections arising from tractography inaccuracies, we retained only those structural connections that were consistently present in at least 90% of participants and we removed cortico-cerebellar spurious connections. A group-level average fiber-length matrix (FL) was then computed, representing the mean streamline length between each reliably connected pair of regions.

Following the approach of [23], we categorized structural connections into short-, middle-, and long-range classes based on the empirical distribution of tract lengths. Specifically, we calculated the 25th percentile (Q1) and 75th percentile (Q2) of the group-level fiber length histogram (see Fig. S4A) and used these values as thresholds:

- Short-range connections: $FL < Q1$
- Middle-range connections: $Q1 \leq FL \leq Q2$
- Long-range connections: $FL > Q2$

These thresholds were subsequently applied to each participant's structural fiber length matrix to classify individual connections accordingly. This analysis included 142 participants, as fiber-length estimation failed for one subject.

Next, we quantified the prevalence of each fiber-length class within the subject-specific DAGs across energetic ranges. For each DAG, we computed the proportion of directed connections falling into the short-, middle-, and long-range categories. Finally, we statistically compared these proportions across energetic ranges to test whether distinct energy levels preferentially support information hierarchical transfer through specific fiber-length classes. Comparisons of the resulting distributions were performed using a Friedman post hoc test for repeated measures ($\alpha = 0.01$).

#### Supplementary Results

To delve deeper into the role of the pattern of firing identified by each dominant eigenvector in shaping the corresponding network $S$ patterns depicted in Fig. 2, we conducted a correlation analysis between the magnitude and phase of each eigenvector and the corresponding $S$ entry.

First, it is noteworthy to highlight the striking spatial patterns exhibited by the principal eigenvector across the three energetic ranges in Fig. S1B, which was extracted by combining subject-level dominant eigenvectors through an analogous complex vectorized sum (after re-shifting by the mean subject phase deviation). In the first range, the spatial firing pattern appears seemingly random, dispersed across various regions. However, in the second range, the ROI-wise vector components demonstrate a notable synchronization tendency, revealing a consistent phase-shifting order aligned with the RSN identity, as indicated by the network-color coding. This synchronizing effect becomes even more pronounced in the last range, where a clear differentiation emerges between sensory-attentive ROIs (i.e., SOM or SVAN) and higher-level transmodal ROIs (i.e., CON).

As illustrated in panel D, progressing from the first to the third energetic range enhances the coupling between spatial and temporal structure: if the phase of node $k$ precedes that of node $j$ (as inferred from the dominant complex eigenmode), it becomes increasingly likely that $S_{jk} > 0$, consistent with directional information flow governed by the antisymmetric nature of the solenoidal matrix. This relationship holds as long as the phase difference remains moderate; $S$ approaches zero when the phase difference nears $\pm\pi$. This phase-dependent organization reinforces the interpretation of $S$ as capturing coherent wave propagation, where sources and sinks are dynamically allocated across the brain's spatial layout.

Furthermore, as $S$ reflects a covariance-based estimate of asymmetric interactions, similar trends are observed when analyzing the co-fluctuation amplitude between nodes $j$ and $k$ $m_{jk}$ —that is, when the product of their signal magnitudes is high, $S_{jk}$ module tends to be more positive in line with dominant phase ordering (Fig. S1C). This alignment between spatial asymmetry, amplitude modulation, and complex eigenmode phase structure reinforces the view that the solenoidal component encodes not just non-reversible interactions, but also the spatiotemporal geometry of traveling-wave-like brain dynamics that grow more pronounced as the system approaches critical slowing down.

More specifically, at R1, amplitude moderately explained *S* (0.35), while phase shifts showed only weak correlation (0.12). This is consistent with phase differences being narrowly distributed around zero, which reduces their ability to explain variance in *S*. In the mid-frequency range (R2), both measures dropped (0.23 and 0.15), reflecting a transitional regime where neither amplitude bursts nor stable phase relationships dominated. In R3, both correlations increased substantially, with phase (0.66) slightly exceeding amplitude (0.60). In this range, the number of effective modes was very low compared to R1 and R2, making the relationship between *S* and the dominant eigenmode more precise and likely contributing to the strong observed correlations.

Analogous trends were observed in Run2 (Fig. S11).

### Supplementary Figures

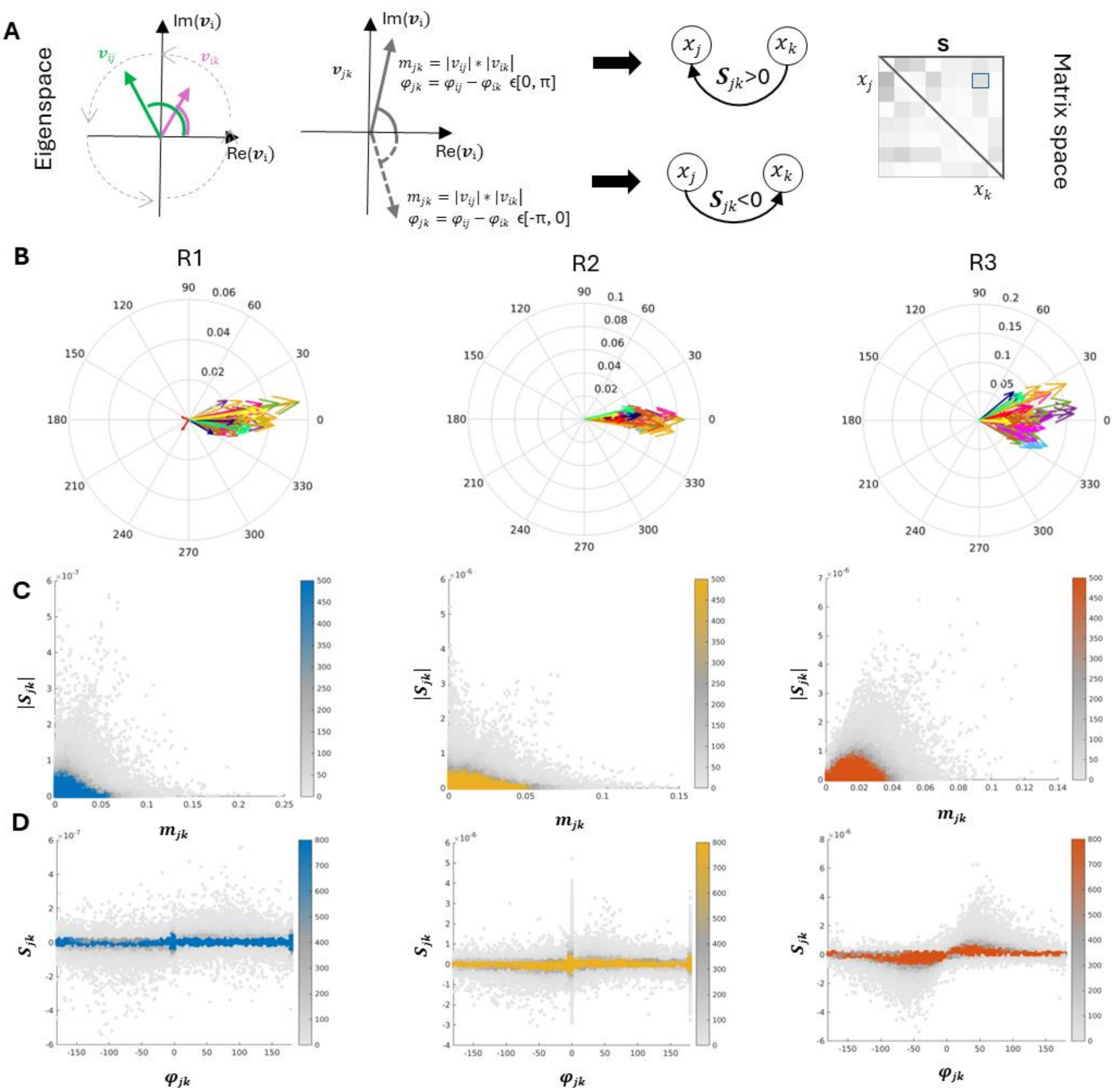


**Figure S1.**

**A)** To demonstrate the mathematical relationship between the eigenspace of the EC and the S-matrix (schematized in panel A), we proceeded as follows: for each subject and frequency range, we extracted the dominant eigenvector (corresponding to the eigenvalue with the least negative decay rate). For each pair of ROIs, the entries of this eigenvector $v_{ik}$ and $v_{ij}$ were compared with the corresponding dc-Cov entry in the S-matrix. By construction, the relationship is coherent: if $S_{jk} > 0$ (and $S_{kj} < 0$, by skew-symmetry, node $k$ drives node $j$, and thus $v_{ik}$ precedes $v_{ij}$ in the rotating plane. Conversely, if $S_{jk} < 0$ (and $S_{kj} > 0$), node $j$ drives node $k$, and $v_{ik}$ follows $v_{ij}$. From this, we defined a new vector

$v_{jk}$ with magnitude $m_{jk}$ (always positive) and phase $\varphi_{jk}$ (positive if $k$ precedes $j$, negative otherwise). We expected $m_{jk}$ to scale with $|S_{jk}|$ and $\varphi_{jk}$ to reflect the signed value of $S_{jk}$.

**B)** group-level dominant eigenvector across the three ranges for Run1. For each range, the across-subject mean eigenvector was computed after realigning individual eigenvectors by their average phase shift. Arrows are color-coded according to large-scale networks (see legend in Fig. 3). Notably, from R1 to R2 and R3, ROIs belonging to the same network tend to cluster together, reflecting large-scale network communication.

**C)** Scatter density plots reporting the correlations described in panel A between the magnitude $|S_{jk}|$ and $m_{jk}$ across subjects for Run1, with mean correlation values of 0.35 (R1), 0.24 (R2), and 0.65 (R3).

**D)** Scatter density plots reporting the correlations described in panel A between the magnitude $S_{jk}$ and $\varphi_{jk}$ for Run1, with mean correlation values of 0.13 (R1), 0.16 (R2), and 0.67 (R3).

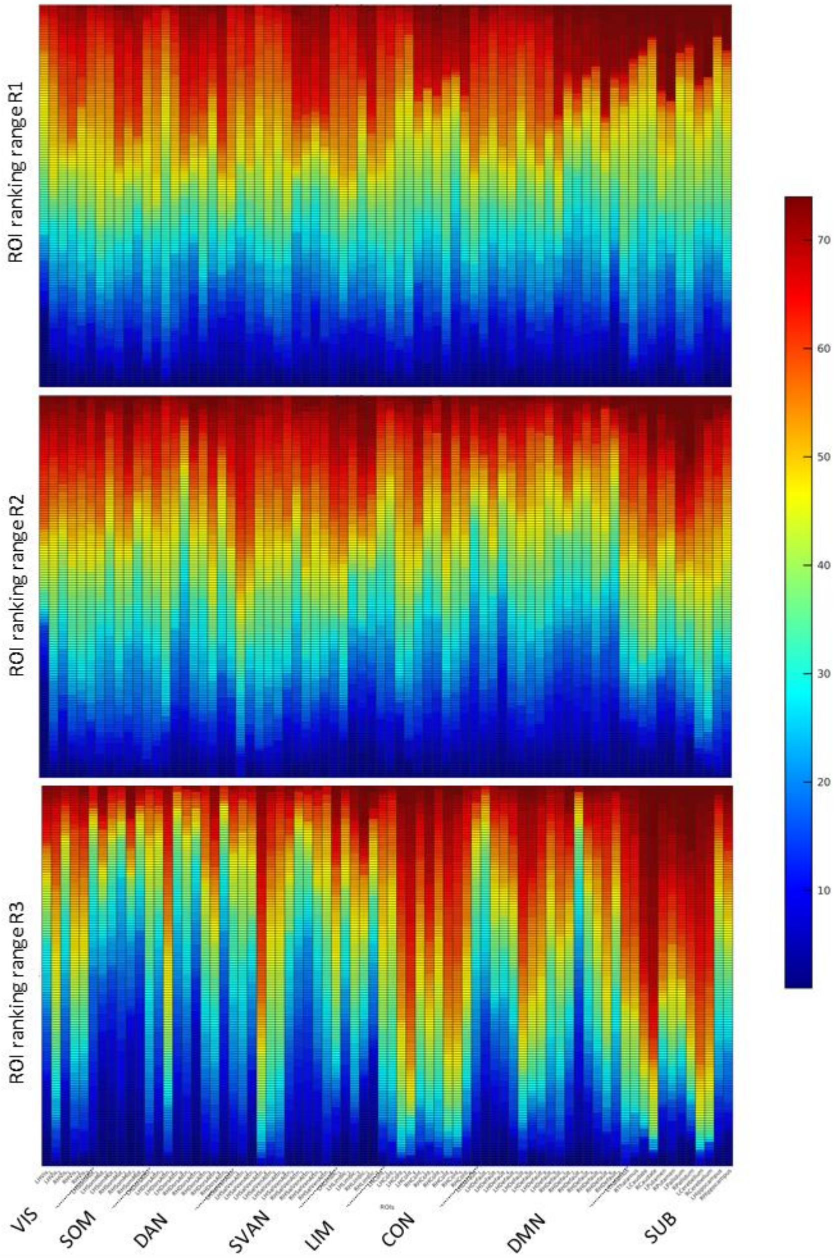


**Figure S2.**

Heatmaps showing the distribution of ROI-wise rankings across subjects, ordered from low to high values, for each of the three ranges (R1–R3) in Run1. As the range progresses from R1 to R3, ROI specialization becomes more pronounced, with individual ROIs spanning progressively narrower ranking intervals. ROIs are grouped by cortical and subcortical networks.

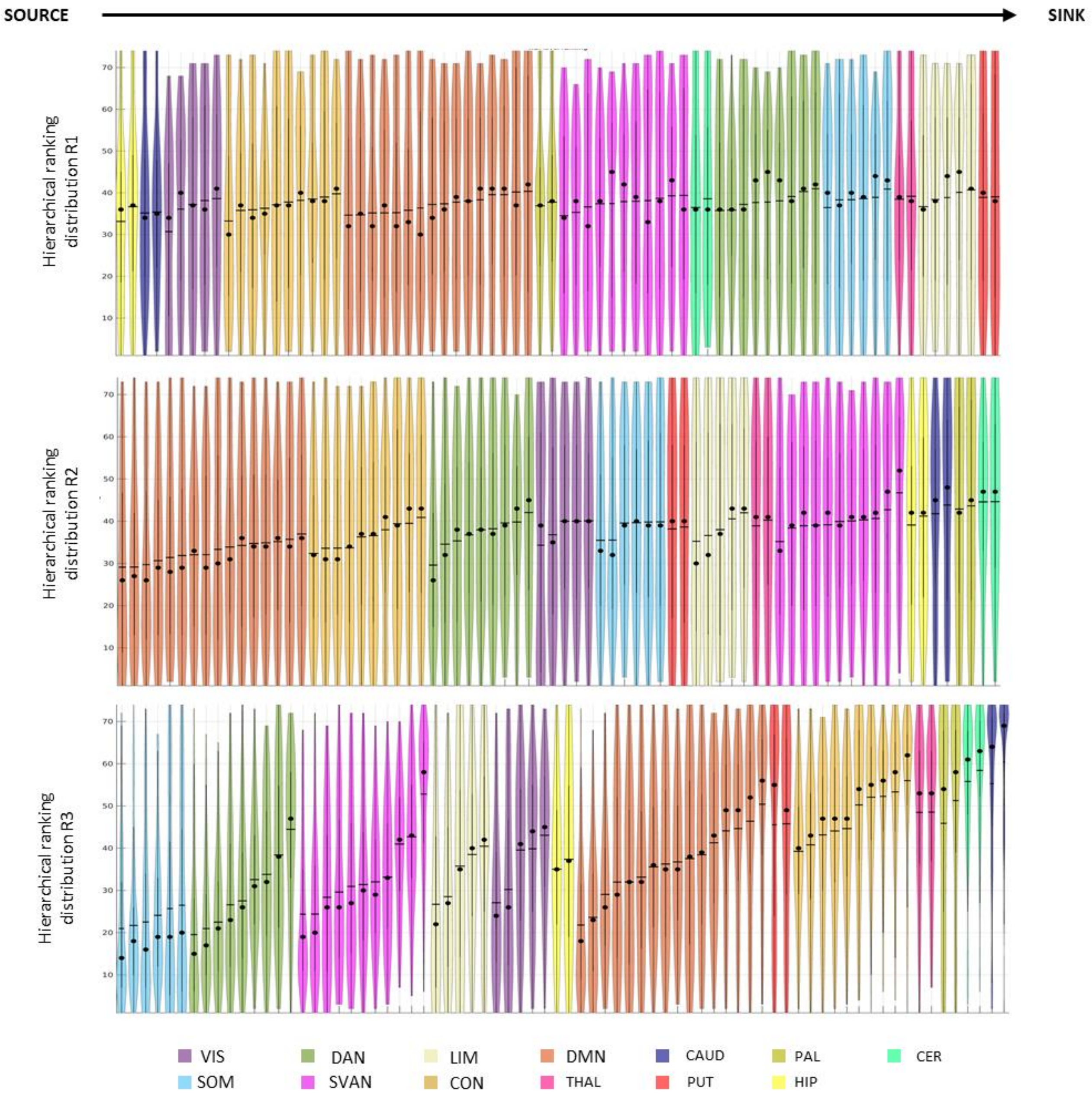


**Figure S3**

Violin plots showing the distribution of hierarchical rankings across subjects and ROIs in the three ranges (R1–R3) for Run1. Brain ROIs are grouped by RSN or subcortical affiliation. Plots are ordered first by the network mean (ascending) and then, within each network subgroup, by ascending ROI values. Horizontal lines mark mean values, and black dots indicate median values.

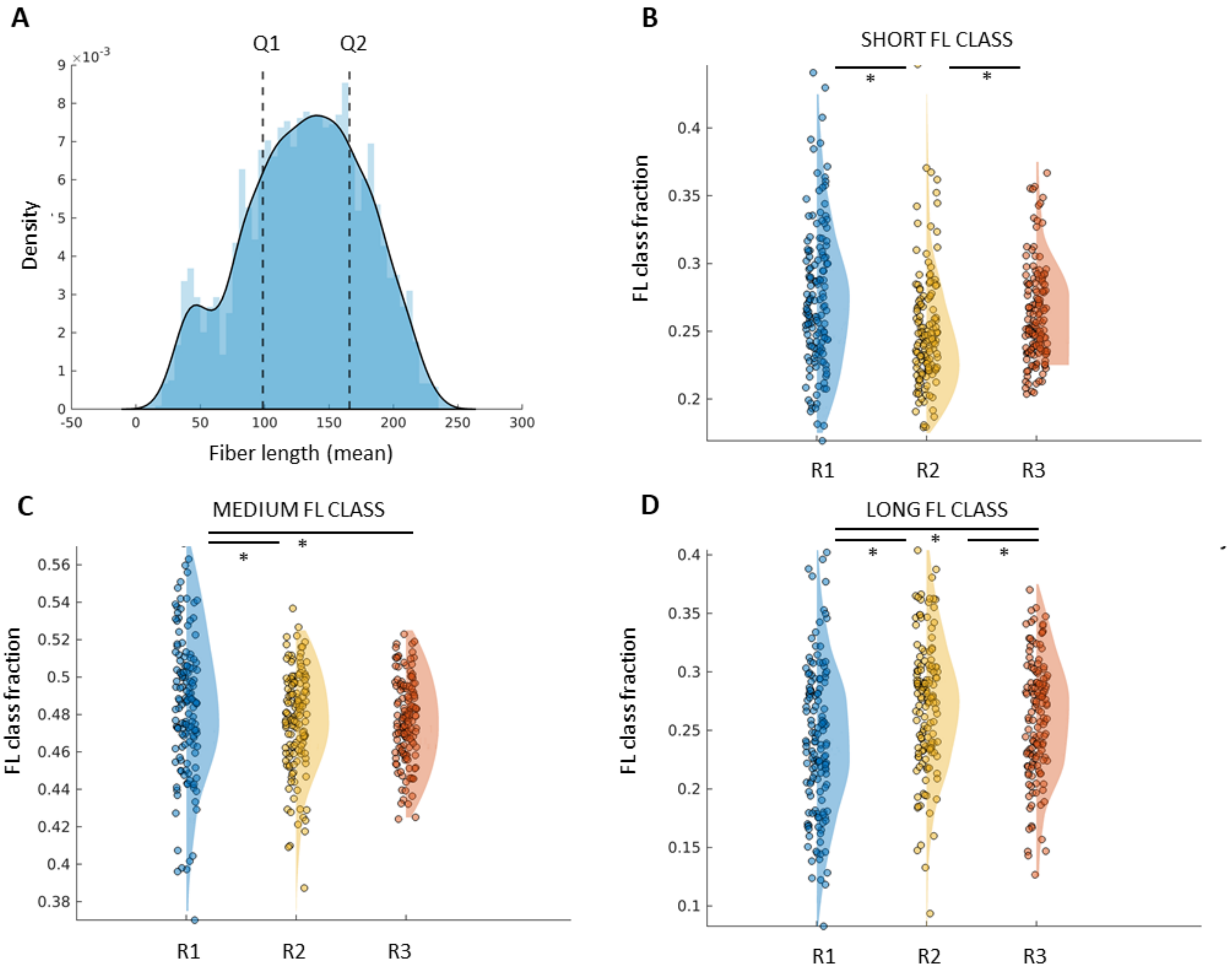


**Figure S4**

- **A)** Distribution of fiber length (FL) values extracted from the group-level FL matrix, as described in the Supplementary Methods. Q1 and Q2 correspond to the $25^{th}$ and $75^{th}$ percentiles, respectively.
- **B)** Group-level comparison of the fraction (%) of connections in the short FL class (<Q1) across ranges for Run1. Significant differences are marked with * (Friedman test with multiple-comparison correction, $\alpha = 0.01$).
- **C)** Group-level comparison of the fraction (%) of connections in the medium FL class (Q1–Q2) across ranges for Run1. Significant differences are marked with *.

**D)** Group-level comparison of the fraction (%) of connections in the long FL class (>Q2) across ranges for Run1. Significant differences are marked with *.

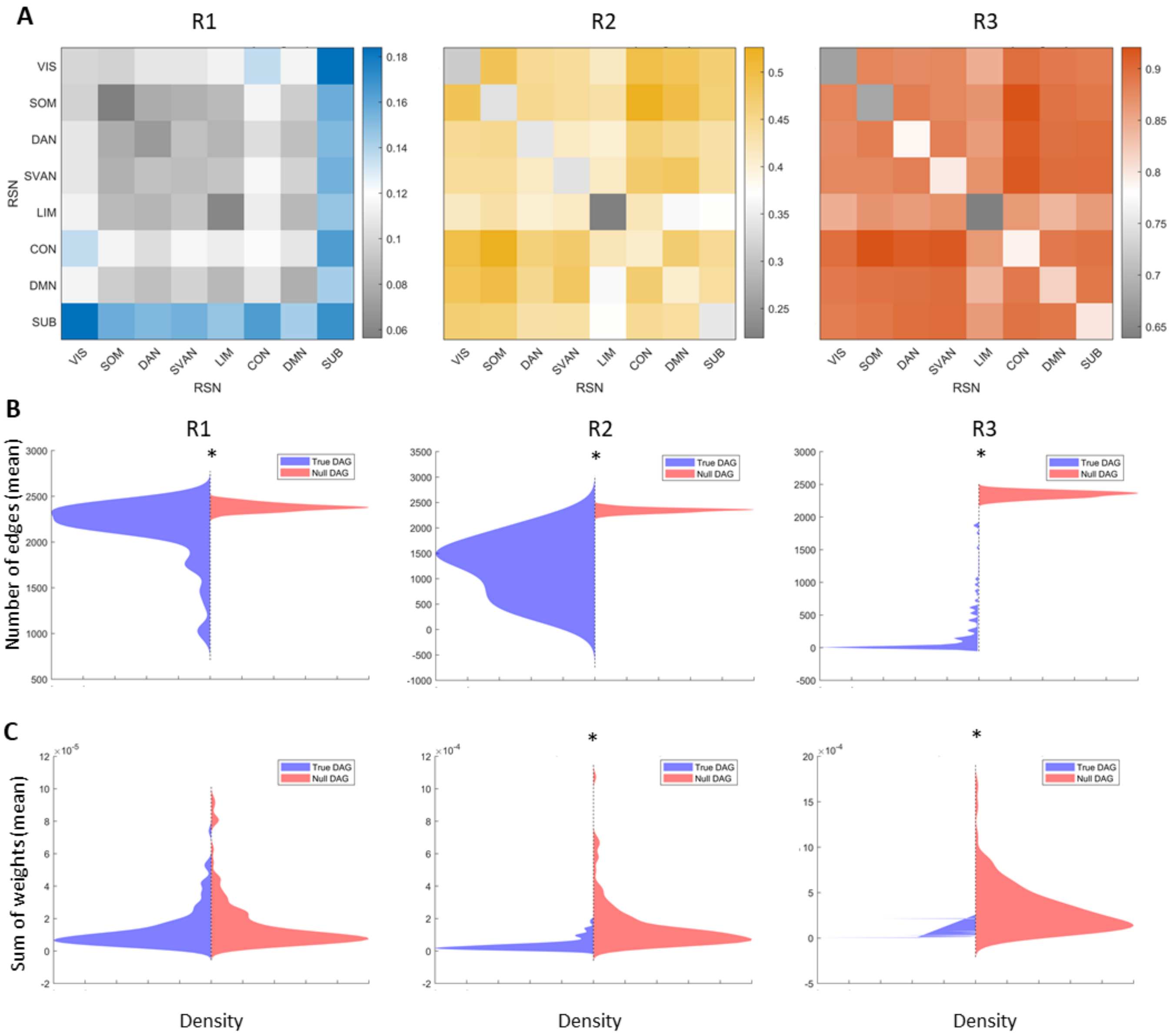


**Figure S5**

A) RSN-level block matrices showing the frequency of link selection in individual DAGs (fraction of total) for each of the three ranges in Run1. It is noteworthy that the frequency values increase from R1 to R3, indicating an increasing stability in the DAG structure across individuals.

B) Comparison of the number of removed edges (mean across 100 iterations) between real and null DAGs across subjects in Run1. Significant differences are indicated by * (Wilcoxon rank-sum test; α = 0.05, corrected for multiple comparisons).

C) Comparison of the sum of weights of removed edges (mean across 100 iterations) between real and null DAGs across subjects in Run1. Significant differences are indicated by * (Wilcoxon rank-sum test; α = 0.05, corrected for multiple comparisons).

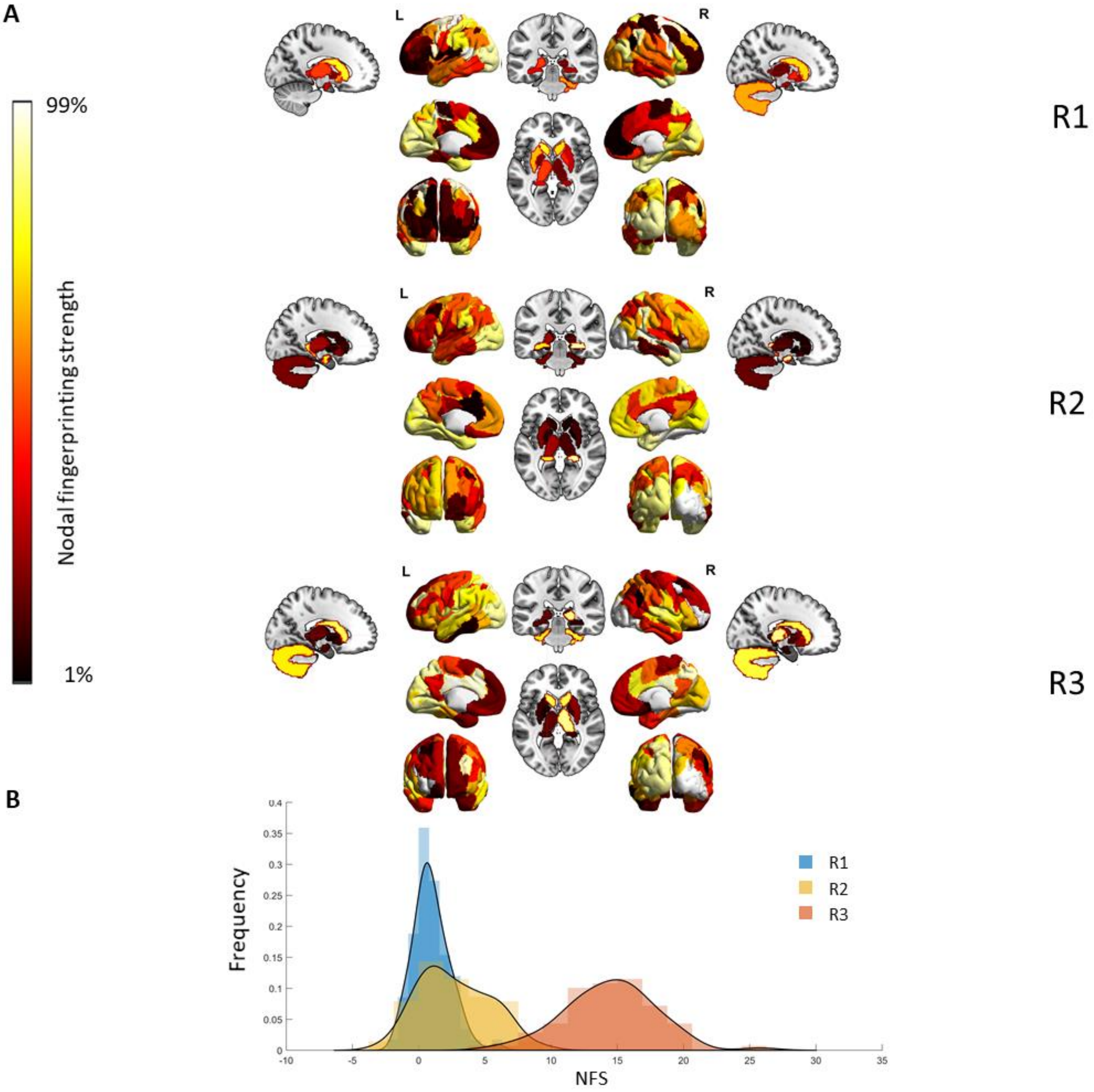


**Figure S6**

A) Brain render showing nodal fingerprinting patterns: ICC-based subject identifiability is represented as nodal fingerprinting strength for each region across the three ranges. Nodal strength was computed as the sum of columns of the ICC edgewise matrix and thresholded at the 1st–99th percentile.

B) Distribution of nodal fingerprinting strength across the three ranges.

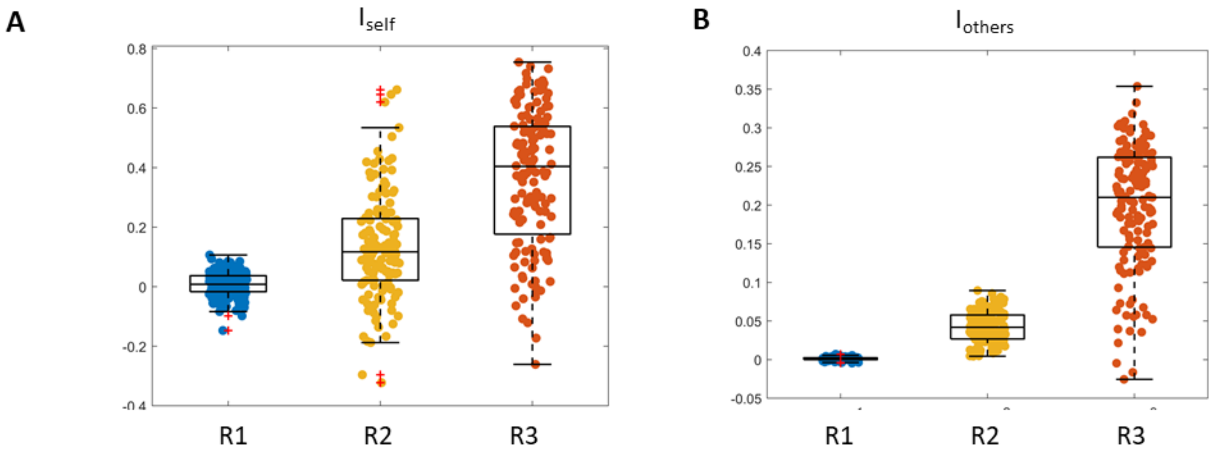


**Figure S7**

Distributions of $I_{self}$, $I_{others}$ across the three ranges. Boxes indicate the mean, standard deviation, and 95% confidence interval of each distribution, with jittered individual data points overlaid.

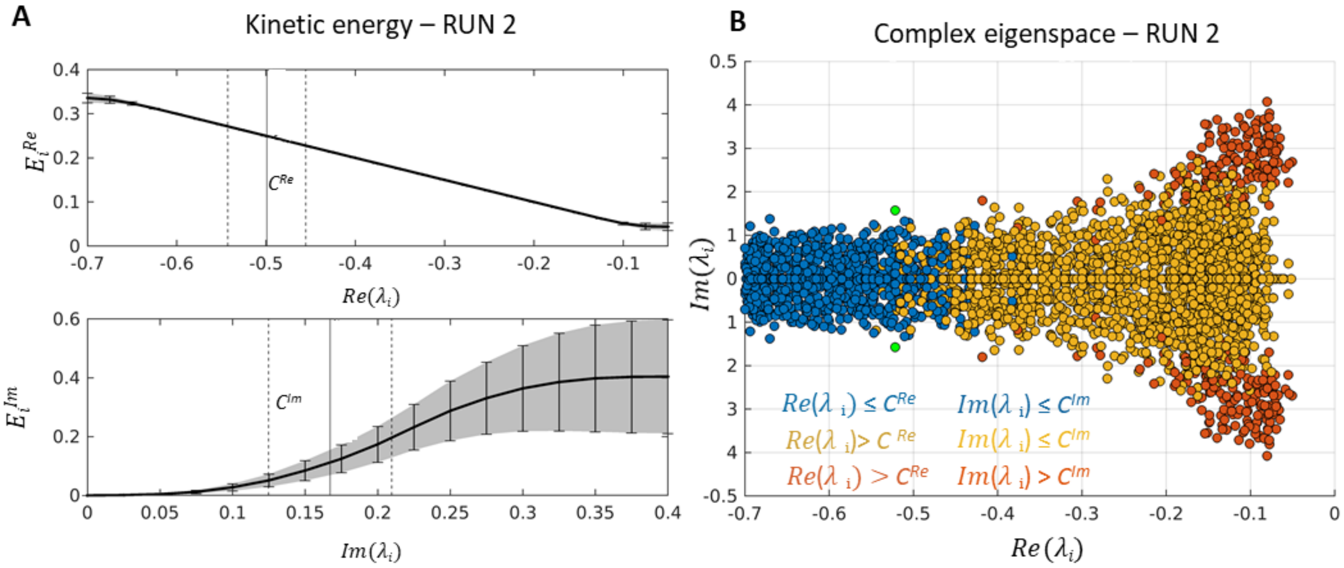


**Figure S8.**

**A)** Group-level plots of the real (top) and imaginary (bottom) components of kinetic energy across the eigenspectrum (solid line = mean, shaded area = standard deviation) for Run2. Cut-offs are indicated (solid line = mean, dashed line = CIs). Because the number of modes varied across subjects, individual curves were interpolated within the displayed range to generate the group representation.

**B)** Scatterplot of complex eigenvalues across all subjects for Run2, shown in the complex plane. As scales progress from left to right, real eigenvalues converge toward zero. Colors indicate energetic range: blue = first, yellow = second, red = third. On average for Run2, R1 encompasses 7.24 modes, R2 15.87 and R3 2.33. Here the fourth range survives just for one subject)

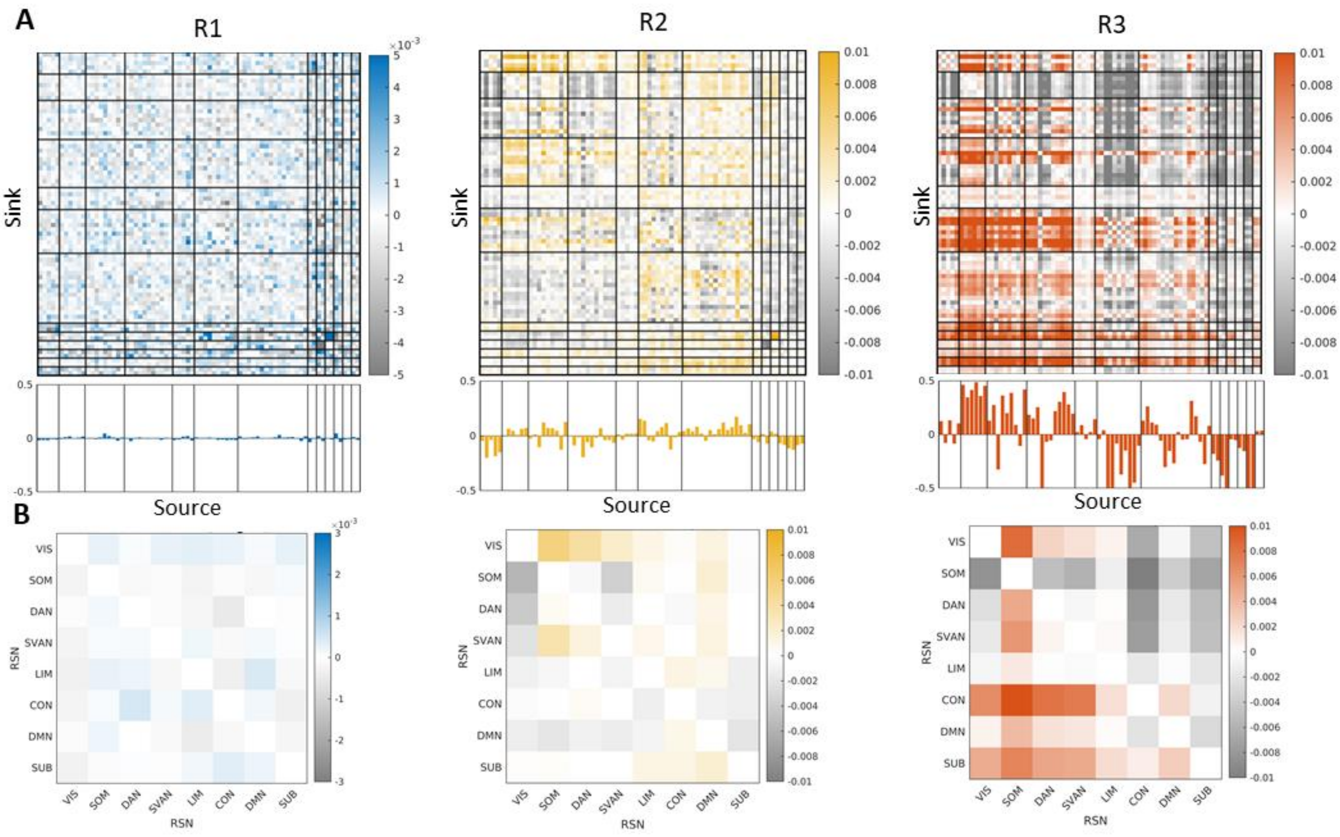


**Figure S9.**

A**)** Group-level average patterns of the differential cross-covariance matrix for the first second and third ranges in Run2. Each individual matrix is normalized by its Frobenius norm, and ROIs are grouped according to cortical and subcortical networks (black lines). Below each matrix, the column-wise strengths are reported: a negative column sum indicates that the node predominantly acts as a receiver, while a positive sum indicates that it predominantly acts as a sender.

**B)** Same matrices as in (A), but averaged across resting-state network (RSN) blocks.

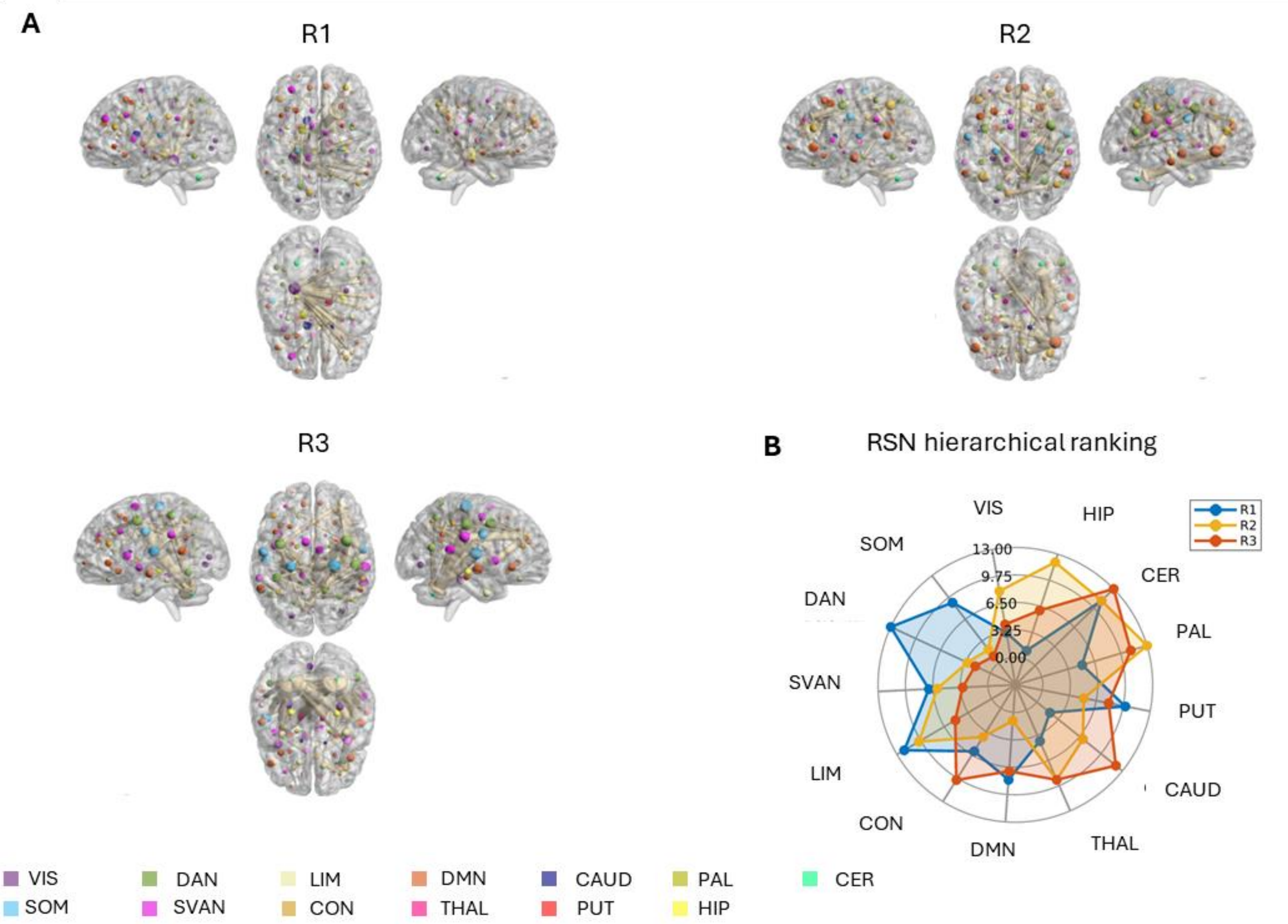


**Figure S10.**
**A)** Mean ROI rankings, derived from each ROI's ranking distribution are shown on a brain rendering to elucidate the spatial distribution of source and target nodes for Run2. Node size reflects the role of the node as a source (larger nodes indicate earlier positions in the hierarchy). Links are displayed if their frequency exceeds the 99th percentile of the DAG frequency matrices (see Supplementary Fig. S15A), with edge thickness proportional to frequency. Node colors correspond to functional networks.
**B)** Spider plot showing mean rankings at the cortical and subcortical network level across ranges (lower values = sources; higher values = sinks).

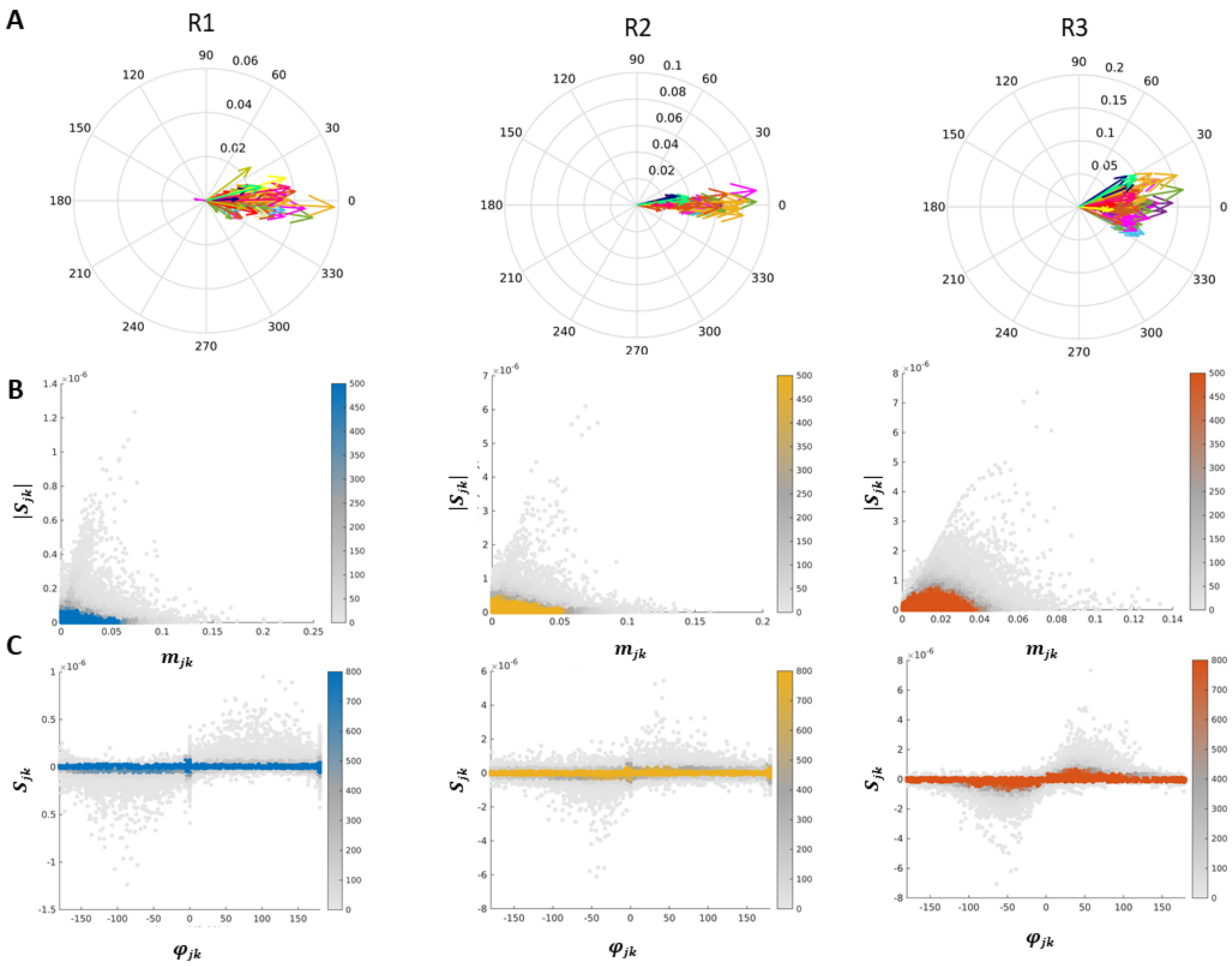


**Figure S11.**

A) Group-level dominant eigenvector across the three ranges for Run2. For each range, the across-subject mean eigenvector was computed after realigning individual eigenvectors by their average phase shift. Arrows are color-coded according to large-scale networks (see legend in Fig. 3). Notably, from R1 to R2 and R3, ROIs belonging to the same network tend to cluster together, reflecting large-scale network communication.

B) Scatter density plots reporting the correlations between the magnitude $|S_{jk}|$ and $m_{jk}$ across subjects, with mean correlation values of 0.36 (R1), 0.19 (R2), and 0.64 (R3).

C) Scatter density plots reporting the correlations between the magnitude $S_{jk}$ and $\varphi_{jk}$, with mean correlation values of 0.15 (R1), 0.12 (R2), and 0.66 (R3).

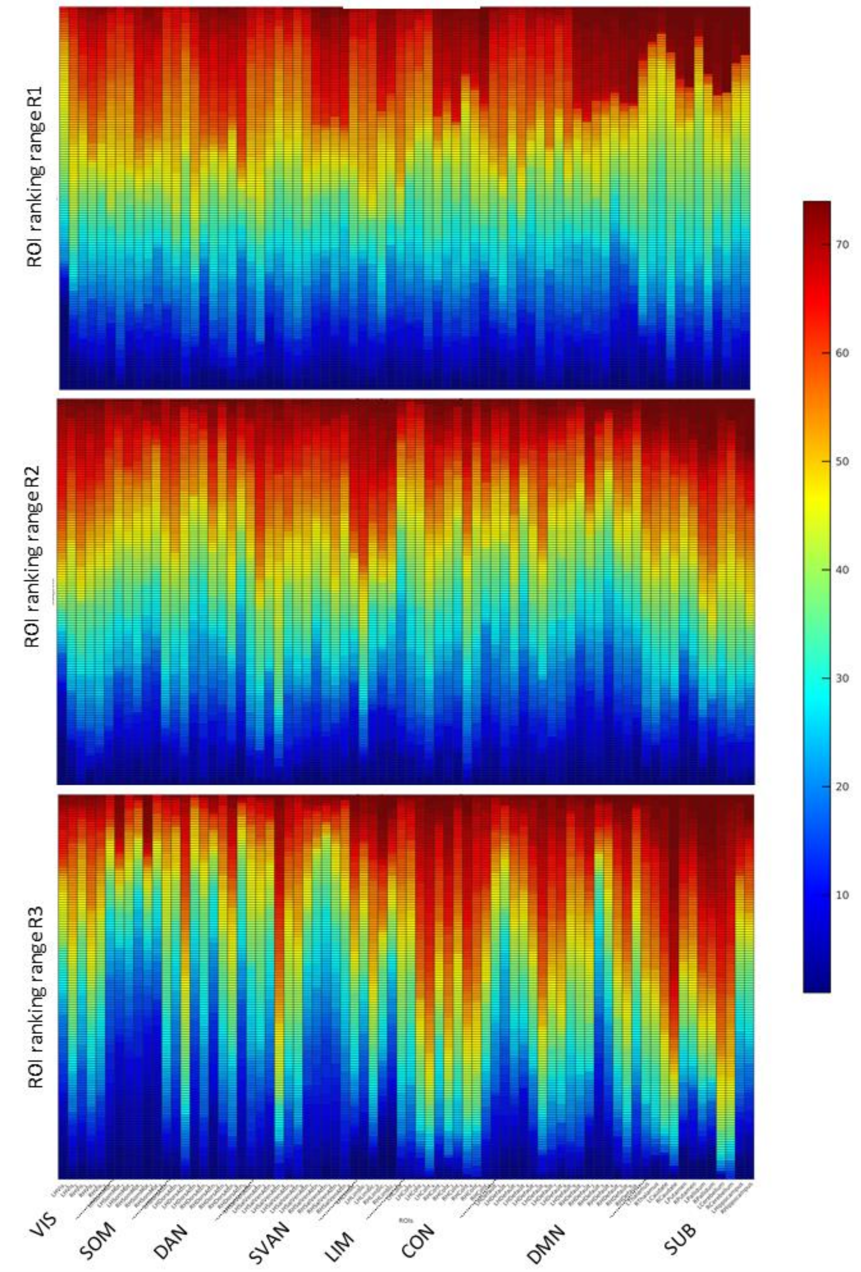


**Figure S12.**

Heatmaps showing the distribution of ROI-wise rankings across subjects, ordered from low to high values, for each of the three ranges (R1–R3) in Run2. As the range progresses from R1 to R3, ROI specialization becomes more pronounced, with individual ROIs spanning progressively narrower ranking intervals. ROIs are grouped by cortical and subcortical networks.

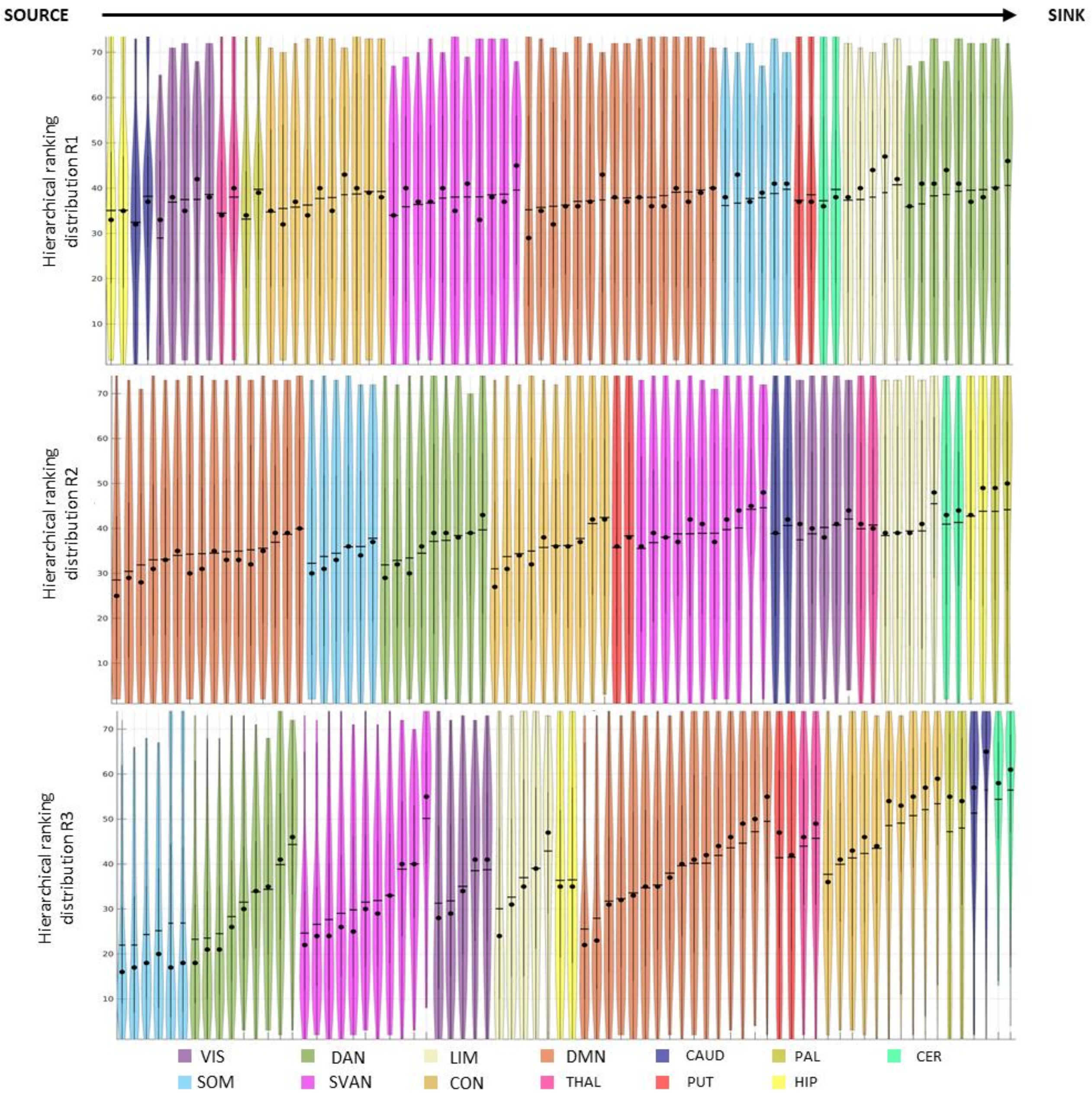


**Figure S13.**

Violin plots showing the distribution of hierarchical rankings across subjects and ROIs in the three ranges (R1–R3) for Run2. Brain ROIs are grouped by RSN or subcortical affiliation. Plots are ordered first by the network mean (ascending) and then, within each network subgroup, by ascending ROI values. Horizontal lines mark mean values, and black dots indicate median values.

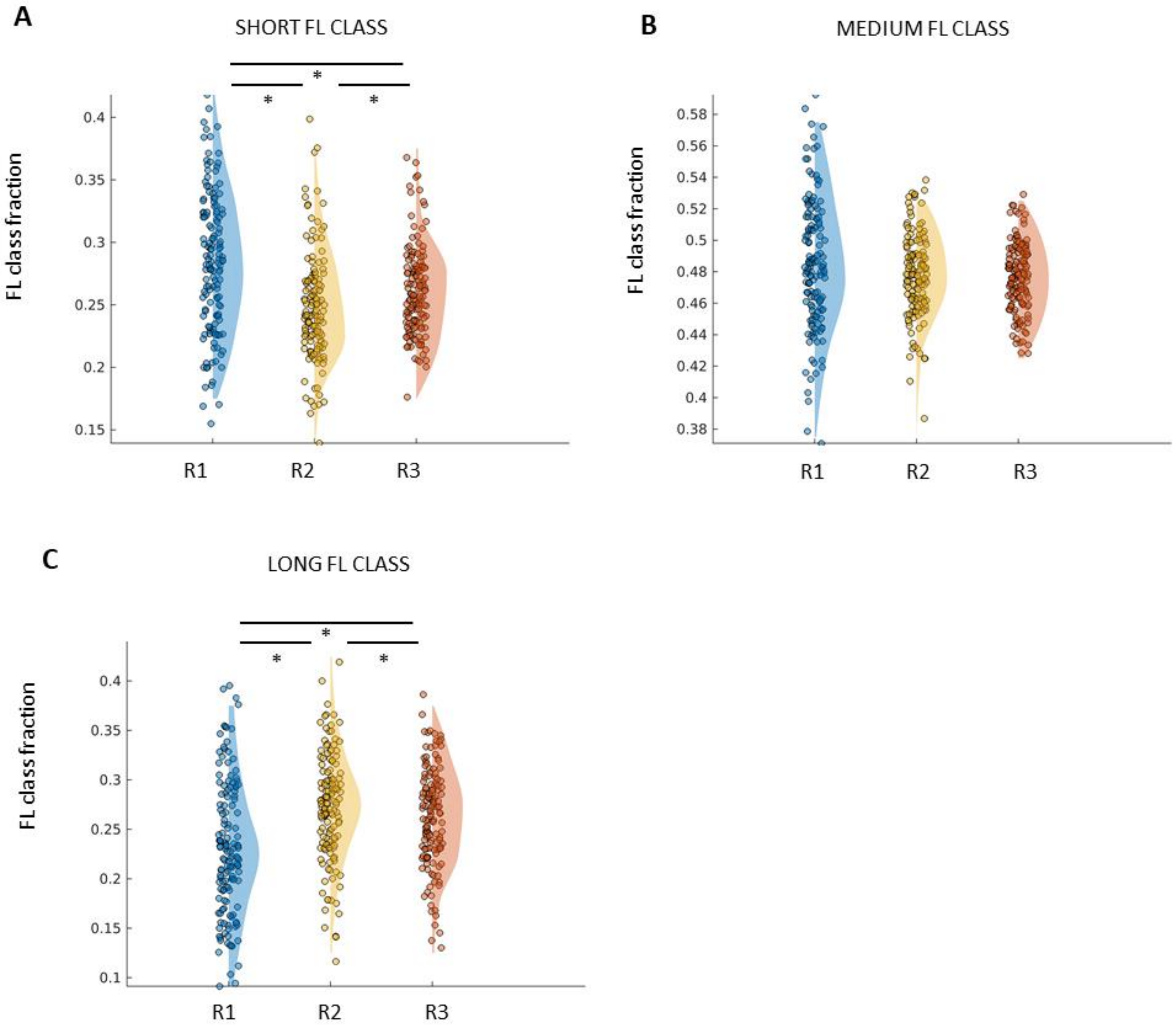


**Figure S14.**

A) Group-level comparison of the fraction (%) of connections in the short FL class (<Q1) across ranges for Run2. Significant differences are marked with * (Friedman test with multiple-comparison correction, $\alpha = 0.01$).

B) Group-level comparison of the fraction (%) of connections in the medium FL class (Q1–Q2) across ranges for Run2. Significant differences are marked with *.

C) Group-level comparison of the fraction (%) of connections in the long FL class (>Q2) across ranges for Run2. Significant differences are marked with *.

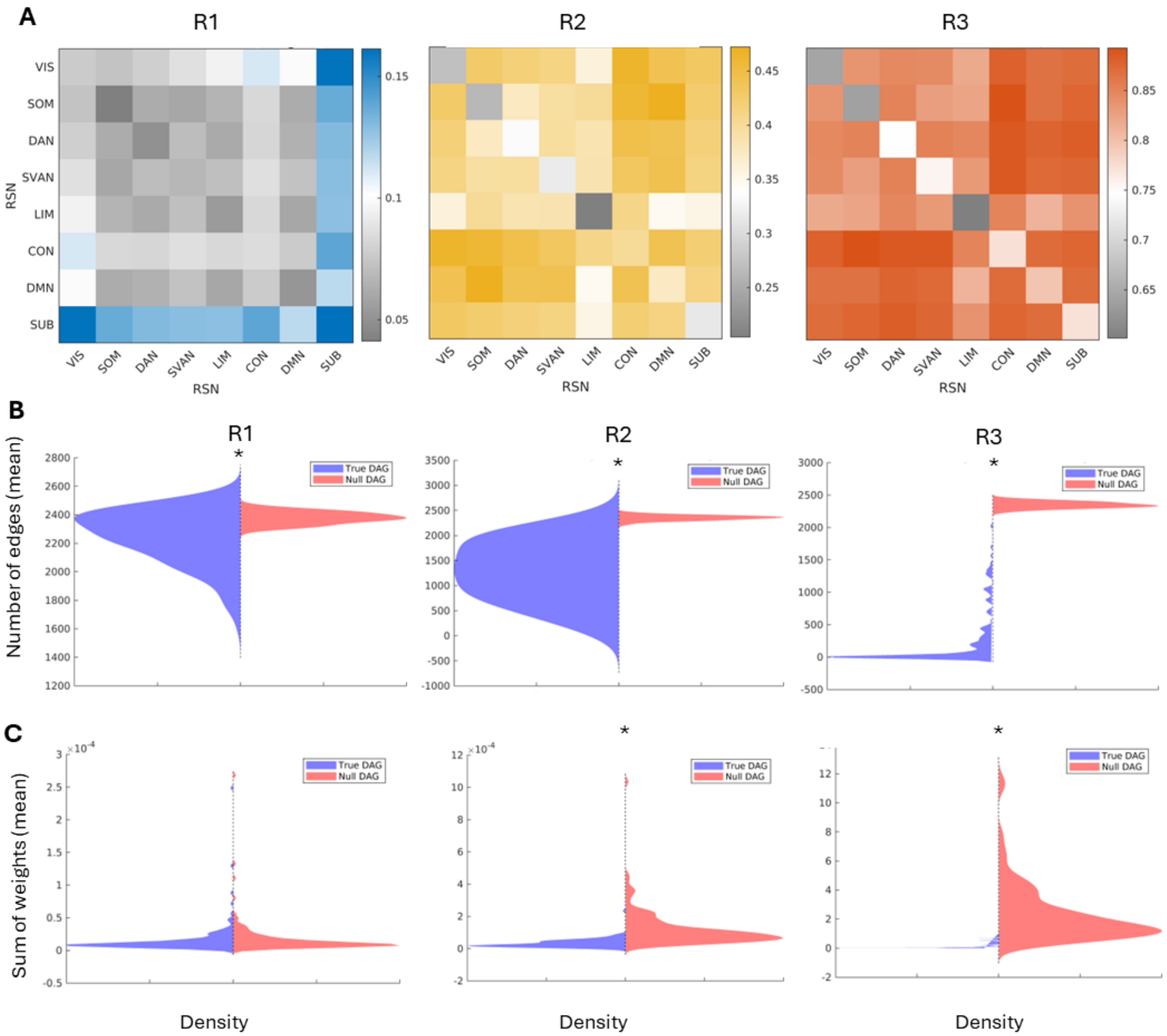


**Figure S15.**

A) RSN-level block matrices showing the frequency of link selection in individual DAGs (fraction of total) for each of the three ranges in Run2. It is noteworthy that the frequency values increase from R1 to R3, indicating an increasing stability in the DAG structure across individuals.

B) Comparison of the number of removed edges (mean across 100 iterations) between real and null DAGs across subjects for Run2. Significant differences are indicated by * (Wilcoxon rank-sum test; $\alpha = 0.05$, corrected for multiple comparisons).

C) Comparison of the sum of weights of removed edges (mean across 100 iterations) between real and null DAGs across subjects for Run2. Significant differences are indicated by * (Wilcoxon rank-sum test; α = 0.05, corrected for multiple comparisons).

#### References


1. Van Essen, D. C. *et al.* The Human Connectome Project: a data acquisition perspective. *NeuroImage* **62**, 2222–2231 (2012).
2. Glasser, M. F. *et al.* The minimal preprocessing pipelines for the Human Connectome Project. *NeuroImage* **80**, 105–124 (2013).
3. Schaefer, A. *et al.* Local-Global Parcellation of the Human Cerebral Cortex from Intrinsic Functional Connectivity MRI. *Cereb. Cortex N. Y. N 1991* **28**, 3095–3114 (2018).
4. Yeo, B. T. T. *et al.* The organization of the human cerebral cortex estimated by intrinsic functional connectivity. *J. Neurophysiol.* **106**, 1125–1165 (2011).
5. Prando, G. *et al.* Sparse DCM for whole-brain effective connectivity from resting-state fMRI data. *NeuroImage* **208**, 116367 (2020).
6. Baron, G., Silvestri, E., Benozzo, D., Chiuso, A. & Bertoldo, A. Revealing the Spatial Pattern of Brain Hemodynamic Sensitivity to Healthy Aging through Sparse Dynamic Causal Model. *J. Neurosci.* **45**, e1940232024 (2025).
7. Ryali, S., Chen, T., Padmanabhan, A., Cai, W. & Menon, V. Development and validation of consensus clustering-based framework for brain segmentation using resting fMRI. *J. Neurosci. Methods* **240**, 128–140 (2015).
8. Taylor, A. J., Kim, J. H. & Ress, D. Characterization of the hemodynamic response function across the majority of human cerebral cortex. *NeuroImage* **173**, 322–331 (2018).
9. Chen, X. *et al.* Leading basic modes of spontaneous activity drive individual functional connectivity organization in the resting human brain. *Commun. Biol.* **6**, 892 (2023).
10. Friston, K. J., Harrison, L. & Penny, W. Dynamic causal modelling. *NeuroImage* **19**, 1273–1302 (2003).
11. Buxton, R. B., Wong, E. C. & Frank, L. R. Dynamics of blood flow and oxygenation changes during brain activation: the balloon model. *Magn. Reson. Med.* **39**, 855–864 (1998).
12. Deco, G., Kringelbach, M. L., Jirsa, V. K. & Ritter, P. The dynamics of resting fluctuations in the brain: metastability and its dynamical cortical core. *Sci. Rep.* **7**, 3095 (2017).
13. Friston, K. J. *et al.* Parcels and particles: Markov blankets in the brain. *Netw. Neurosci.* **5**, 211–251 (2021).
14. Benozzo, D. *et al.* Analyzing asymmetry in brain hierarchies with a linear state-space model of resting-state fMRI data. *Netw. Neurosci.* 1–42 (2024) doi:10.1162/netn_a_00381.
15. Casti, U. *et al.* Dynamic Brain Networks with Prescribed Functional Connectivity. (2023).

16. Liégeois, R., Santos, A., Matta, V., Van De Ville, D. & Sayed, A. H. Revisiting correlation-based functional connectivity and its relationship with structural connectivity. *Netw. Neurosci. Camb. Mass* **4**, 1235–1251 (2020).
17. Friston, K. J. *et al.* Dynamic causal modelling revisited. *NeuroImage* **199**, 730–744 (2019).
18. Lin, T. W., Das, A., Krishnan, G. P., Bazhenov, M. & Sejnowski, T. J. Differential Covariance: A New Class of Methods to Estimate Sparse Connectivity from Neural Recordings. **2733**, 2709–2733 (2018).
19. Jeurissen, B. & Szczepankiewicz, F. Multi-tissue spherical deconvolution of tensor-valued diffusion MRI. *NeuroImage* **245**, 118717 (2021).
20. Smith, R., Skoch, A., Bajada, C., Caspers, S. & Connelly, A. *Hybrid Surface-Volume Segmentation for Improved Anatomically-Constrained Tractography*. (2020).
21. Smith, R. E., Tournier, J.-D., Calamante, F. & Connelly, A. Anatomically-constrained tractography: Improved diffusion MRI streamlines tractography through effective use of anatomical information. *NeuroImage* **62**, 1924–1938 (2012).
22. Smith, R. E., Tournier, J.-D., Calamante, F. & Connelly, A. SIFT2: Enabling dense quantitative assessment of brain white matter connectivity using streamlines tractography. *NeuroImage* **119**, 338–351 (2015).
23. Meijer, K. A., Steenwijk, M. D., Douw, L., Schoonheim, M. M. & Geurts, J. J. G. Long-range connections are more severely damaged and relevant for cognition in multiple sclerosis. *Brain* **143**, 150–160 (2020).
